# Identification and comparison of somatic antigen composition for bacteria from *Providencia* genus

**DOI:** 10.1101/2022.04.05.486866

**Authors:** Polina D. Kuchur, Anna I. Churkina, Anna A. Rybina, Aleksey S. Komissarov

## Abstract

*Providencia* is a genus of Gram-negative bacteria belonging to the Morganellaceae family. This genus includes nine species (P. *stuartii, P. sneebia, P. rettgeri, P. rustigianii, P. heimbachae, P. burhodogranariea, P. alcalifaciens, P. huaxiensis*, and *P. vermicola*) with varying degrees of virulence, capable of infecting humans and insects [1, 2].

For Gram-negative bacteria, the somatic antigen (O-antigen) has become one of the key virulence factors. It is the highly immunogenic part of lipopolysaccharides due to the distal location. O-antigens are characterized by structural heterogeneity, providing varying degrees of inter- and intraspecific virulence. At the genetic level, somatic antigens have an operon structure. Operon genes responsible for the synthesis and transformation of O-polysaccharide are transcribed together. Analysis of O-antigen operon organization determines genes specific for each O-serogroup. It is beneficial for molecular typing of strains and for studies of bacterial evolution.

This study focuses on identifying and comparing candidates for O-antigen operons in *Providencia* species with different levels of virulence. The hypothesis is the presence of an association between the O-antigen operon composition and the bacteria lifestyle. Data processing and analysis are carried out by a pipeline developed by the authors. Pipeline combines five steps of the genome analysis: genome quality evaluation, assembly annotation, operon identification with verification of operon boundaries, and visualization of O-antigen operons. The results reveal previously undescribed O-antigen genes and the changes in the O-antigen operons structure. Among the changes are a transposon insertion leading to tetracycline resistance and the presence of IS elements.

## Introduction

*Providencia* is one of the eight genera of the Morganellaceae family, previously classified as Enterobacteriaceae [3]. This genus consists of seven species, which are widespread and include opportunistic human pathogens, insect pathogens, and symbionts [4]. *P. stuartii* and *P. rettgeri* are the most common opportunistic human microorganisms that can cause inflammation of the urinary tract in catheterized patients [5]. These microorganisms, along with *P. rustigianii* and *P. alcalifaciens*, are the cause of gastrointestinal diseases and nosocomial infections in humans [6]. *P. rettgeri, P. stuartii, P. rustigianii*, and *P. alcalifaciens* are commonly found in water and soil. *P. vermicola* was isolated from nematodes [7], and rohu fish [8], *P. sneebia* and *P. burhodogranaeriae* were isolated from *Drosophila melanogaster* [9]. The importance of *P. heimbachae, P. vermicola*, *P. sneebia*, and *P. burhodogranaeriae* in human morbidity has not been studied.

Lipopolysaccharides (LPS) have a significant impact on the pathogenicity of Gram-negative bacteria. LPS are also called endotoxins [10] and are located on the outer membrane of the bacterial cell wall. Glycoconjugate composed of 3 components:

1. lipid A which is the most conserved lipid part of endotoxin;
2. inner and outer core oligosaccharides, connecting lipid A with O-antigen;
3. O-polysaccharide (or O-antigen due to the antigenic specificity) exposed to the environment.

O-antigen is the most variable component of LPS, which performs both structural and immunogenic functions of Gram-negative bacteria. Variations in the O-antigen structure occur due to sugar residue modifications and changes in chemical bonds between oligosaccharides [11]. Changes in the somatic antigen structure determines the potential for the bacterial pathogenicity. Located on the outer membrane of bacteria, the O-antigen is capable of triggering the host’s immune response by activating the complement system [12]. The diversity of the polysaccharide structure makes it possible to bypass the host’s defense mechanisms and colonize new niches [13]. Remarkably, the O-antigen is characterized by both inter- and intraspecies heterogeneity of the structure. Based on this, O-antigen is used for serotyping of Gram-negative bacteria.

At the genetic level the O-antigen has an operon or cluster structure [14]. In these structures, genes are grouped on the same chromosome site and are transcribed together. The O-antigen gene cluster includes three groups of genes [13]:

- genes for biosynthesis of monosaccharides;
- glycosyltransferase genes;
- genes for processing and transfer of O-antigen to the outer membrane of the bacterial cell wall.

Currently, active research is underway in the field of molecular identification and classification of clinically significant *Providencia* species. They are based on the study of the O-antigen structure [15]. Methods of the nuclear magnetic resonance spectroscopy, serological methods and sequencing determined the structures of O-polysaccharides and the serological relationship of three *Providencia* species (*P. alcalifaciens, P. stuartii*, and *P. rustigianii*). Despite slight differences in the composition and size of the O-antigen operon between *Providencia* species, they all contain the main genes for the biosynthesis of O-polysaccharide. There are the *rml* and *gal* genes responsible for the synthesis of monosaccharides, the genes for glycosyltransferases, the O-antigen flippase (*wzx*) and polymerases (*wzy*) [16].

We hypothesize that the O-antigen operon organization is associated with the biological properties of bacteria and their lifestyle. If the hypothesis is confirmed, analysis of the operon composition will improve the classification of bacteria (O-serotyping) and suggest ways of bacterial evolution. This study aims to identify and compare candidate O-antigen operons of *Providencia* species. It includes searching for genes of O-antigen biosynthesis in the literature and creating a custom bioinformatics pipeline. Pipeline should be capable for genome annotation, O-antigens search, visualizing, and comparing O-antigen operons in selected bacterial species.

## Materials and methods

### Data source, data quality assessment, and annotation

All complete *Providencia* genomes available to July 2021 were downloaded from the National Center for Biotechnology Information (NCBI). To study O-antigen operons, we selected complete genome assemblies without plasmids with the N50 value equal to its total sequence length. The last criterion was chosen to exclude genomes with large stretches of ambiguous bases, which might reduce the quality of the operon annotation (for example, disrupting operon integrity). Assemblies with plasmids were removed from further analysis because O-antigen genes can be represented both in the genome and in plasmids [17].

### Pipeline development. Identification and visualization of O-antigen operons

In this research, we developed the pipeline to identify and visualize candidate O-antigen operons in *Providencia* genomes. It includes steps of genome quality evaluation, genome annotation, search for somatic antigen genes, identification of all operons, finding, and visualization of the O-antigen operons. Distinctive features of our pipeline are annotation of genomes with several tools, prediction and validation of operon boundaries, and a step for additional annotation of unknown genes.

The quality of assemblies was assessed using QUAST v5.0.2 [18] with default parameters. All assemblies were annotated with Prokka v1.14.6 [19] with default parameters. PGAP [20] annotations of studied genomes were retrieved from the NCBI RefSeq data FTP site in general feature format (GFF). As an additional annotator, EggNOG v2.0.1 [21] tool was used.

The boundaries of operons were identified via Operon-Mapper [22] with default options. Candidate O-antigen operons were found based on GFF annotation. A list of genes associated with O-polysaccharide biosynthesis from the Ovchinnikova et.al [23] study was used as a reference. Operon numbers, boundaries of operons and GFF annotation were used to search for and visualize candidate O-antigen operons. Custom Python scripts based on DNA Features Viewer library [24] can be found on Github (https://github.com/rybinaanya/O-antigens). Verification of operon boundaries was performed by finding promoters and terminators. For promoters and terminators search BPROM and FindTerm [25] were used. For clarifying gene annotation, we ran BLAST [26] and ORFfinder [27] with default parameters. Genes homologous to O-antigen related genes from the Ovchinnikova et al. study were identified within the O-antigen operon using NsimScan [28] with default parameters.

### Construction of gene phylogenetic trees

The conserved *galE* gene from the *galETKM* operon was common to all *Providencia* species and was used in the phylogenetic analysis. Nucleotide sequences of *galE* gene were aligned via the MAFFT algorithm [29]. A Neighbor-Joining phylogenetic gene tree was built at the MAFFT web service with the number of bootstrap iterations set to 500.

## Results

### Initial data quality

28 genomes of *Providencia* species (Table S1 in Supplementary materials) were analyzed to find candidate O-antigen operons. Four assemblies represented unknown species. The rest belonged to one of the five *Providencia* species (*P. stuartii*, *P. alcalifaciens*, *P. rettgeri, P. heimbachae*, or *P. rustigianii*) (Table 1). All assemblies met the selection criteria.

**Table 1:**
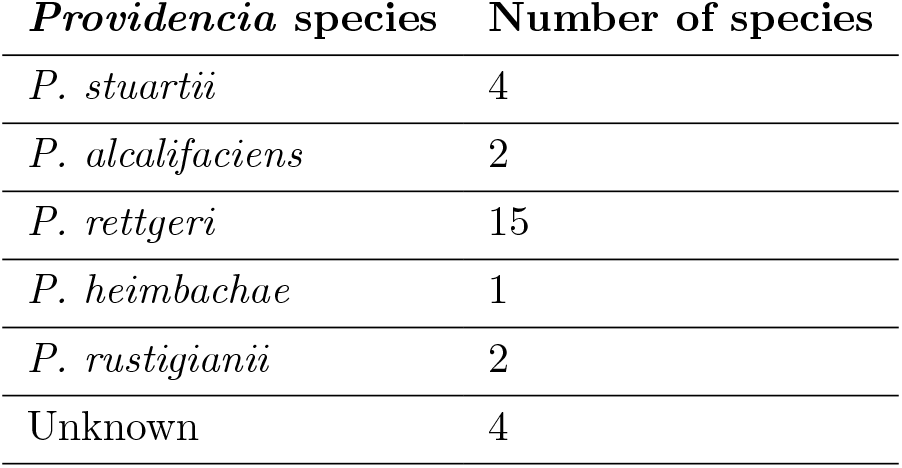
Number of assemblies for each *Providencia* species used in the study.

### Search and visualization of candidate operons for O-antigen

The custom pipeline identified an O-antigen operon similar to that described by Ovchinnikova et al. (Fig. 1, Table S3 in Supplementary materials). Among studied genomes this O-antigen operon differed by number and order of genes. However, its functional composition seems complete meaning the presence of nucleotide sugar biosynthesis genes, genes involved in glycosyl transferring and O-unit formation, and O-antigen processing genes.

**Figure 1.**
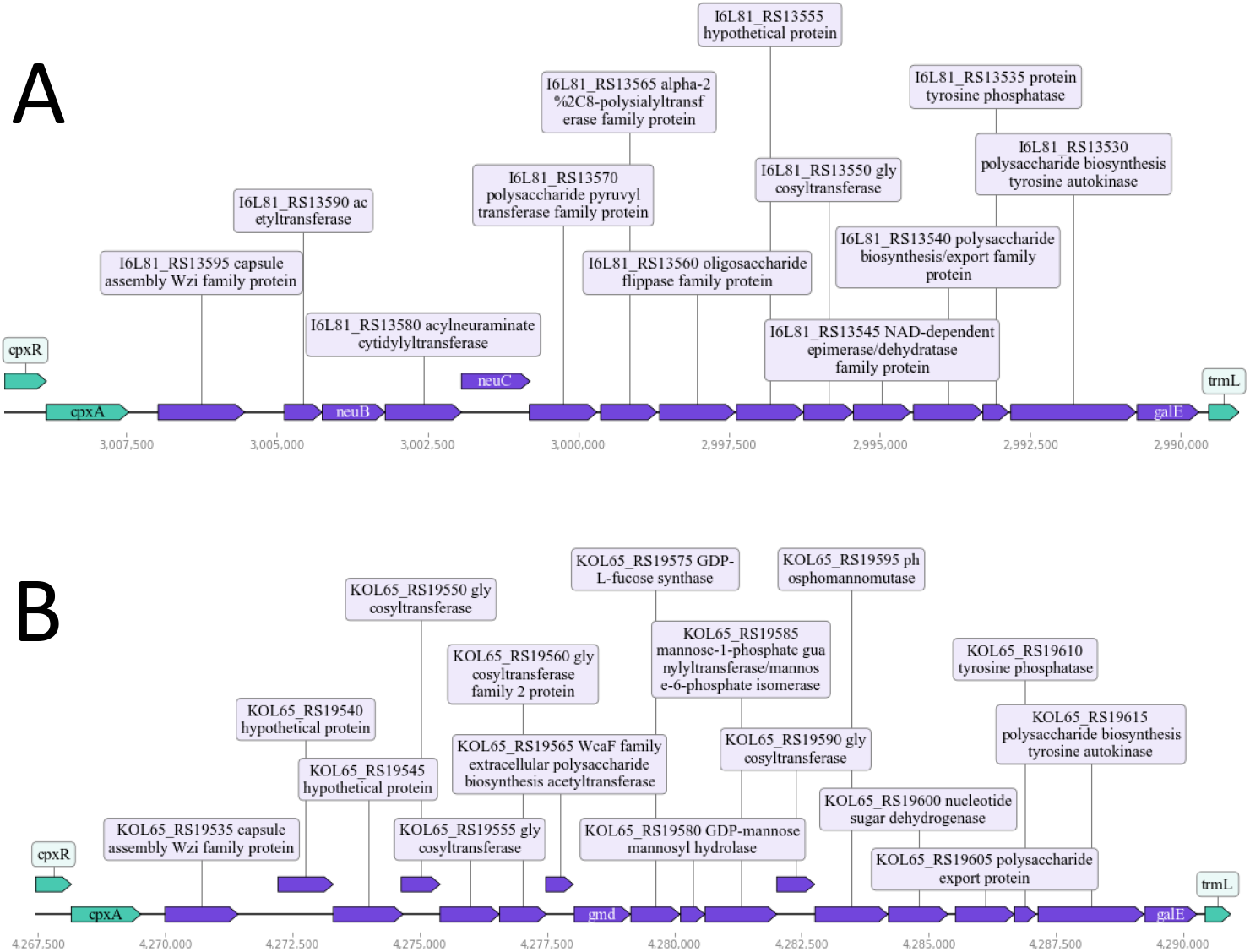
Organization of the O-antigen operon in *P. rettgeri* strain FDAARGOS 1451 (A) and *P. rettgeri* strain PreM15973 (B). The O-antigen operon is colocalized with the *cpxRA* and *wzi* operons. For each gene annotated by PGAP, either gene name or locus tag and product are indicated.

Noteworthy, the O-antigen operon was flanked by the *trmL* gene and the *cpxRA* operon in all studied *Providencia* species. The *cpxRA* operon was adjacent to the *wzi* operon in most of the studied genomes. The exceptions were *P. alcalifaciens* and *P. rettgeri*, where *wzi* was absent. The *cpxR/cpxA* genes encode the two-component system and are associated with virulence features and biofilm formation [30]. The *wzi* gene is associated with the K-antigen synthesis [31]. The proximity of O-antigen- and K-antigen-related genes on the chromosome possibly indicates a tight connection between biosynthesis pathways of O-specific and capsular polysaccharides.

### O-antigen operon with transposon insertion

In several cases, the O-antigen operon contained genes related to antigen biosynthesis as well as genes derived from mobile elements such as transposons and insertion elements (Fig. 2). In *P. rettgeri* strain BML2496, the operon structure was altered due to the transposon incorporation. Transposon was flanked by the IS4 family sequences and carried *tetD*, *tetA*, *tetC*, and *tetR* genes. It enabled *P. rettgeri* strain BML2496 to establish the resistance to tetracycline [32]. *TetD* gene encodes a transcription regulator [33]. *TetA* is responsible for the synthesis of the tetracycline efflux MFS transporter [34]. The *tetC* and *tetR* genes are required for the tetracycline resistance-associated transcriptional repressor TetC and tetracycline resistance transcriptional repressor TetR production [35, 36]. The insertion of transposon led to breakdown of the gene encoding the flippase synthesis. A remarkable feature of this insertion might be the ability to be transcribed in both directions.

**Figure 2.**
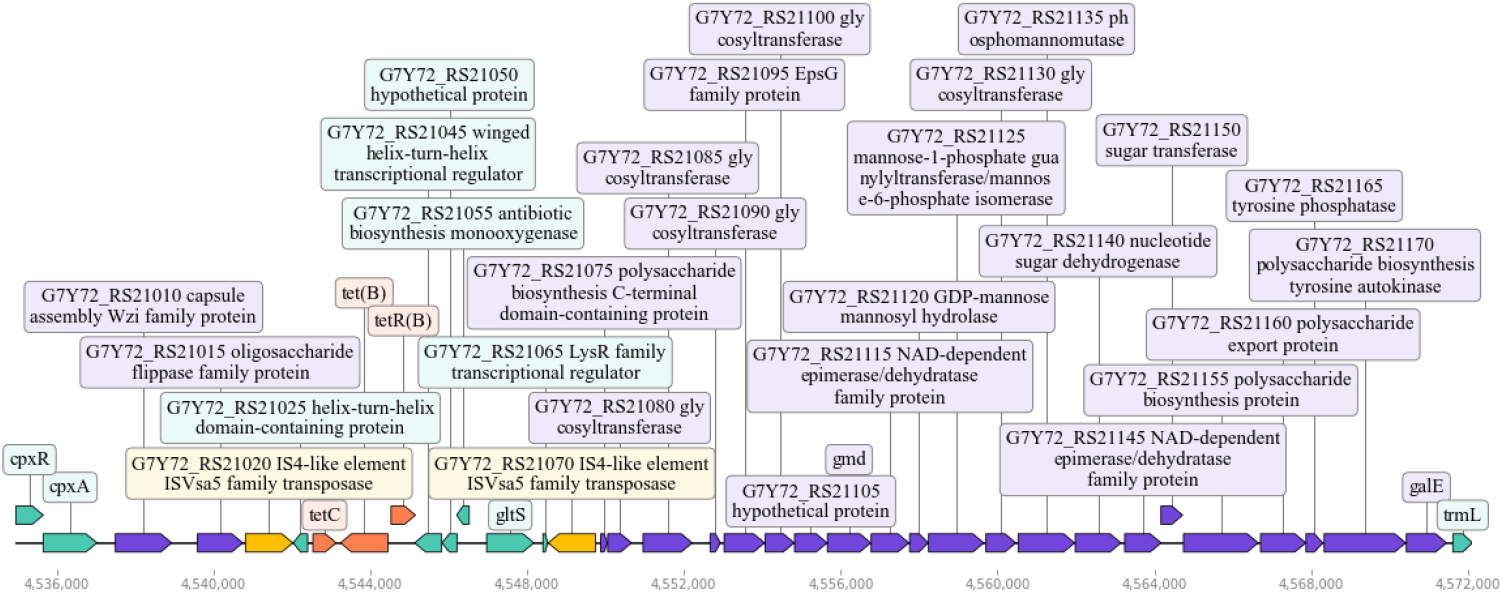
The O-antigen operon in *P. rettgeri* strain BML2496 with a transposon insertion. Antibiotic resistance genes (orange) are surrounded by genes encoding transposases, each annotated as IS4 family transposase ISVsa5 (yellow). For each gene annotated by PGAP, either gene name or locus tag and product are indicated.

Opportunistic human bacteria can cause inflammation of the urinary tract. According to the bacteria lifestyle [37], a possible pathway for the transposon occurrence could have been in response to treatment of urogenital infections with tetracycline [38]. Another acquired gene that can provide bacteria with the antimicrobial activity might be *yqjZ*. Although the function of this gene was not clear, we can assume its involvement in the antibiotic production due to the 100% similarity with the gene ECKY1497_P10770 from *Escherichia coli* KY 1497 according to BLAST. ECKY1497_P10770 gene encodes antibiotic biosynthesis monooxygenase [39].

The length of the O-antigen operon in *P. rettgeri* significantly differed from other *Providencia* species. The average operon length in *P. rettgeri* was 24 kb nucleotides compared to 20 kb nucleotides in other bacteria (Table S2 in Supplementary materials). The transposon insertion was able to considerably increase the distance between genes adjacent to the operon (*cpxR* and *trmL*). The size of transposon in the genome of *P. rettgeri* strain BML2496 was approximately 8 kb.

### O-antigen operons with IS elements

In several O-antigen operons, insertion of mobile genetic element occurred without bringing foreign genes of antimicrobial resistance. These operons are classified here as IS-containing O-antigen operons. We found examples of incorporation of insertion sequence (IS) into the O-antigen operon that caused gene breakdown.

*P. rettgeri* strain PreM15758 had the IS3 family transposase insertion between genes *rfbM* and *algA* (Fig. 3). Based on the KEGG annotation, *rfbM* gene encodes mannose-1-phosphate guanylyltransferase. As the algA gene has the same function [40], we hypothesized that *rfbM* and *algA* could be fragments of some gene disrupted by IS incorporation. We ran BLAST on the fragments surrounding IS which allowed us to suggest the existence of original gene *manC*. The sequence fragment near the transposase (hypothetical protein on Fig. 3) was another transposase of the same family with 95.15% of similarity. The *algA* fragment could be a part of *manC* gene with 98.53% of identity and neighboring hypothetical protein as a glycosyltransferase. These results support our assumption on the breakdown of *manC* in two fragments (*algC* and *rfbM*) because of the IS element appearance.

**Figure 3.**
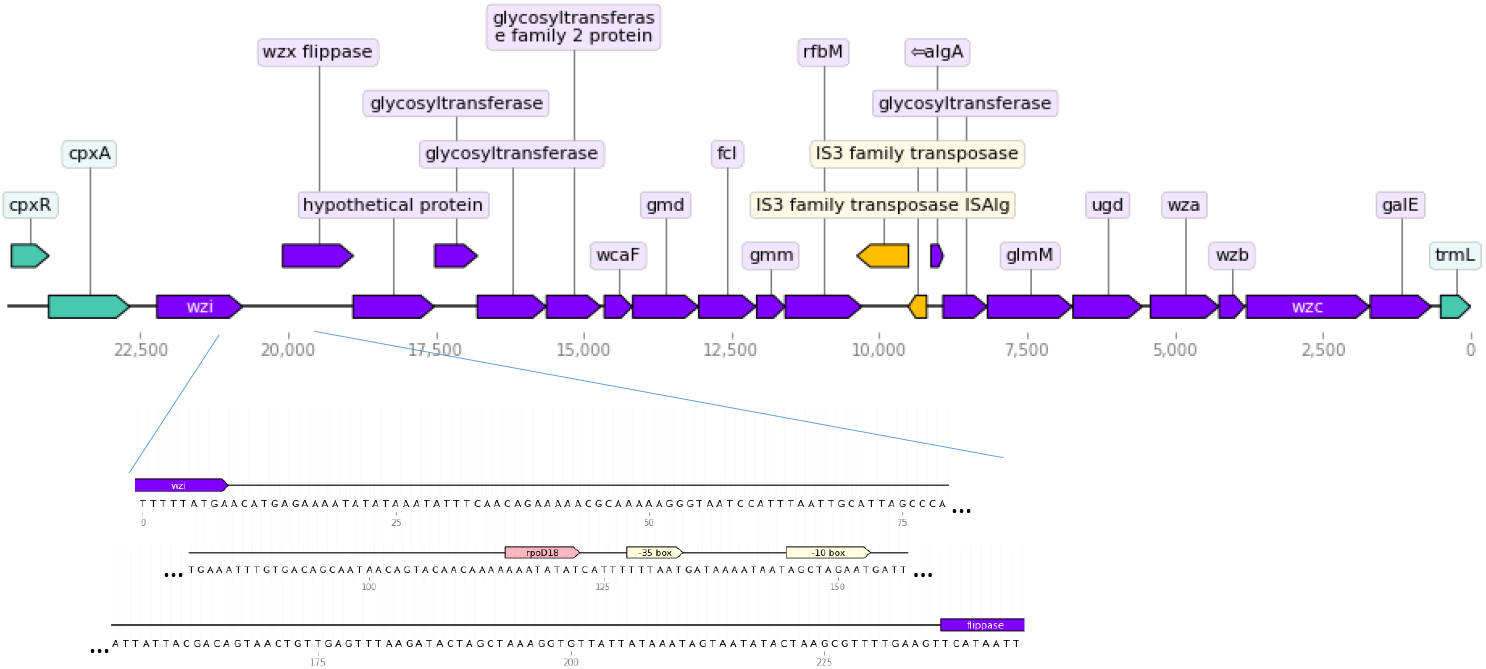
*P. rettgeri* strain PreM15758 O-antigen with the IS elements insertion (yellow). Annotation of genes is based on Prokka.

We suppose the link between O-antigen operon composition with the IS insertion and bacteria lifestyle. O-units formation can be abnormal as IS element disturbed the mannose-1-phosphate guanylyltransferase responsible for transferring phosphorus containing nucleotide groups. According to the organism description [41], it was obtained from the urine of the patient with prostate cancer.

In *P. stuartii* strain AR_0026, the IS element embedding probably resulted in the appearance of four fragments between the gene coding for polysaccharide pyruvyl transferase and *epsG* (Fig. 4). The first fragment annotated as *epsJ* might encode glycosyltransferase with 327 amino acids in length instead of 148 as in the case of *P. stuartii* strain AR 0026. The nucleotide segment in a range from *epsJ* to *epsG* was annotated via BLAST search as a region coding for two fragments of glycosyltransferase family 25 protein (as it was found in unclassified *Proteus*) divided by the IS5 family transposase. Thus, it could be the second example of the gene disruption due to the IS element insertion. Based on the lifestyle, *P. stuartii* strain AR 0026 is characterized as host-associated or clinical pathogen [42]. This feature closes *P. stuartii* to previously described *P. rettgeri* PreM15758 and supports our hypothesis that pathogenic *Providencia* strains tend to have O-antigen operons with genes destroyed by IS elements.

**Figure 4.**
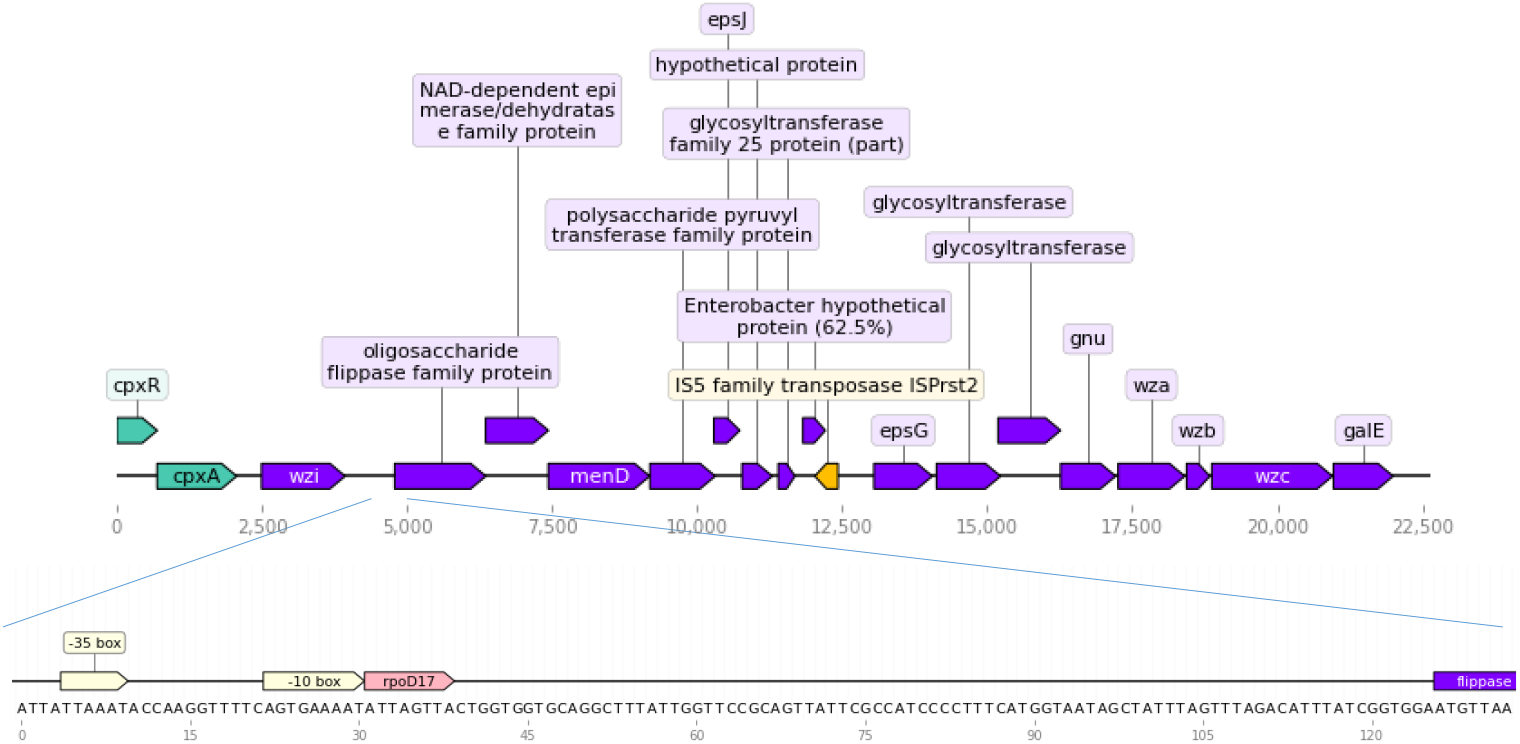
O-antigen composition of the *P. stuartii* strain AR 0026. The IS element is marked yellow. Annotation of genes is based on Prokka.

The size of the O-antigen operon in *P. rustigianii* strain NCTC6933 was estimated to be two times less than in strains above (Fig. 5). In particular, the operon lacked for *wzi, wza, wzb*, and *wzc* genes usually bordering the O-antigen part in other *Providencia*. IS elements were co-directed with other genes in contrast to *P. rettgeri* strain PreM15758 and *P. stuartii* strain AR 0026. Manual annotation of genes upstream and downstream of transposases suggested them encoding glycosyltransferase fragments. Isolation source of *P. rustigianii* strain NCTC6933 has not been reported, so we can not check the hypothesis in this case.

**Figure 5.**
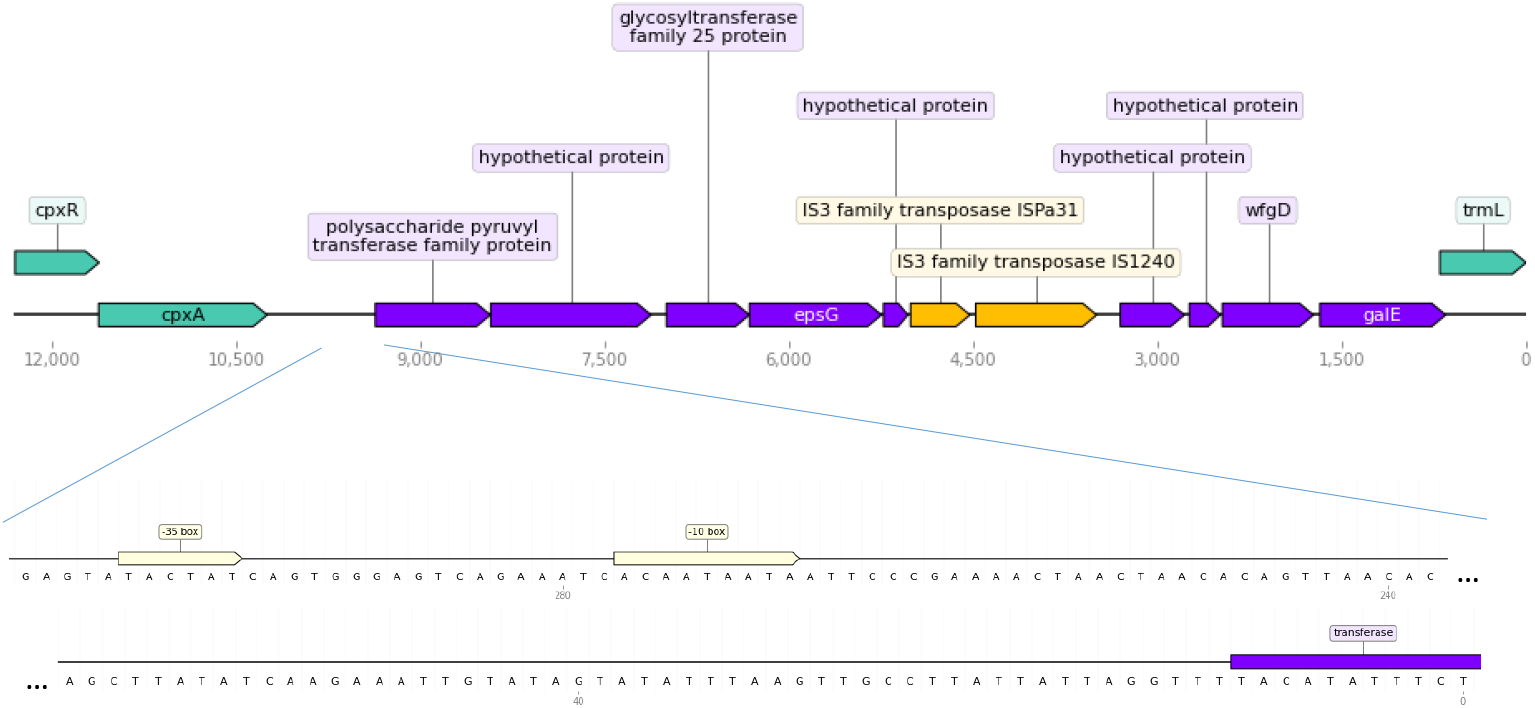
*P. rustigianii* strain NCTC6933 O-antigen composition. The IS element is marked by yellow. Annotation of genes is based on Prokka.

The *P. rustigianii* strain NCTC8113 showed the presence of the IS element in O-antigen operon (Fig. 6). The IS element inserted in a UDP-N-acetylglucosamine 4,6-dehydratase (*wbgZ*) gene, splitting it into two fragments. After the IS element removal, the smaller fragment was confirmed to be a part of the *wbgZ* gene though it was not annotated by Prokka. We could not clarify the nature of the hypothetical protein encoded by gene near *cpxA*. Isolation source of this bacteria has not been reported as for *P. rustigianii* strain NCTC6933.

**Figure 6.**
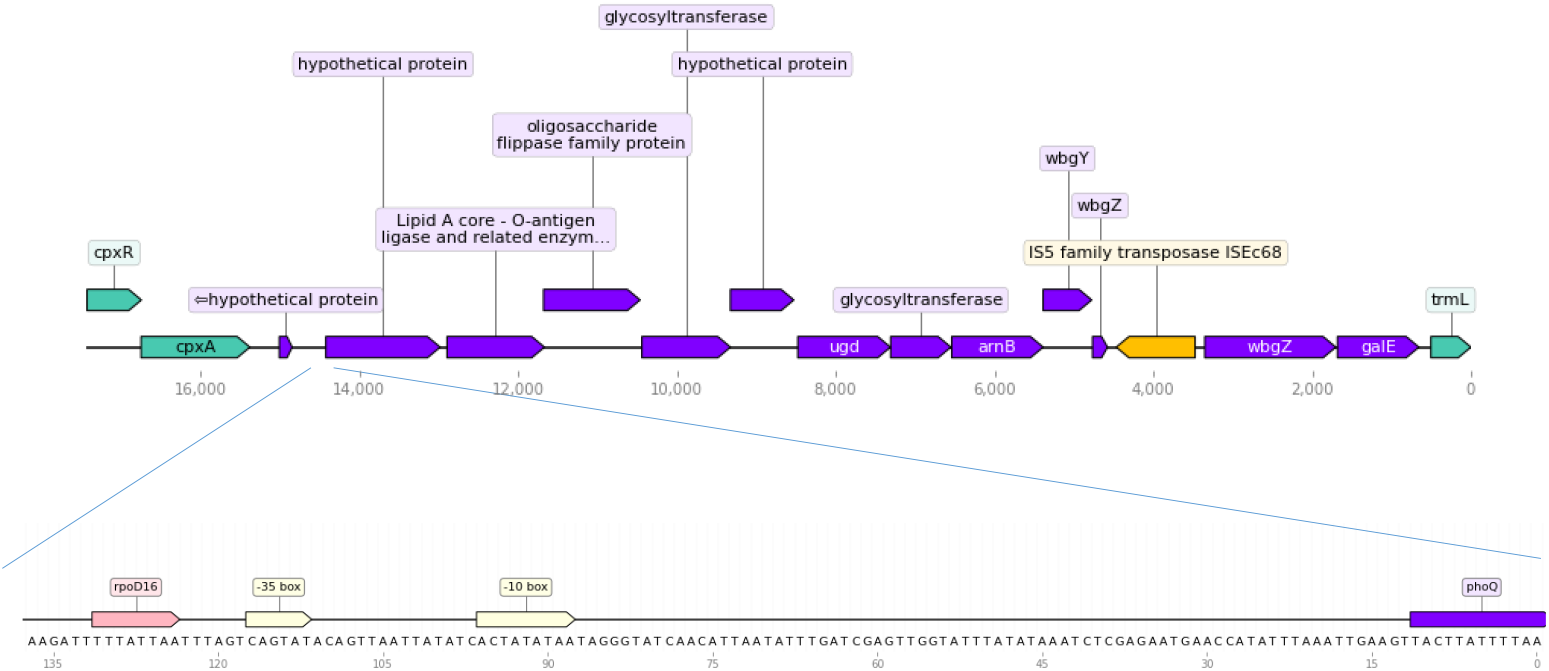
O-antigen composition of the *P. rustigianii* strain NCTC8113. The IS element is marked by yellow. Annotation of genes is based on Prokka.

In *P. heimbachae* strain NCTC12003, transposase insertion destroyed the gene without the possibility of its reconstruction (Fig. 7). The loss of neighboring genes or their parts was not excluded because the gaps between the genes are considerable (up to 1500 base pairs). Along with the insertion element, the glycosyltransferase gene that was annotated only with EggNog, probably obtained nonsense mutation resulting in the appearance of stop codon in the middle of coding sequence. *P. heimbachae* strain NCTC12003 [43] was obtained from penguin feces. The host health state is unknown.

**Figure 7.**
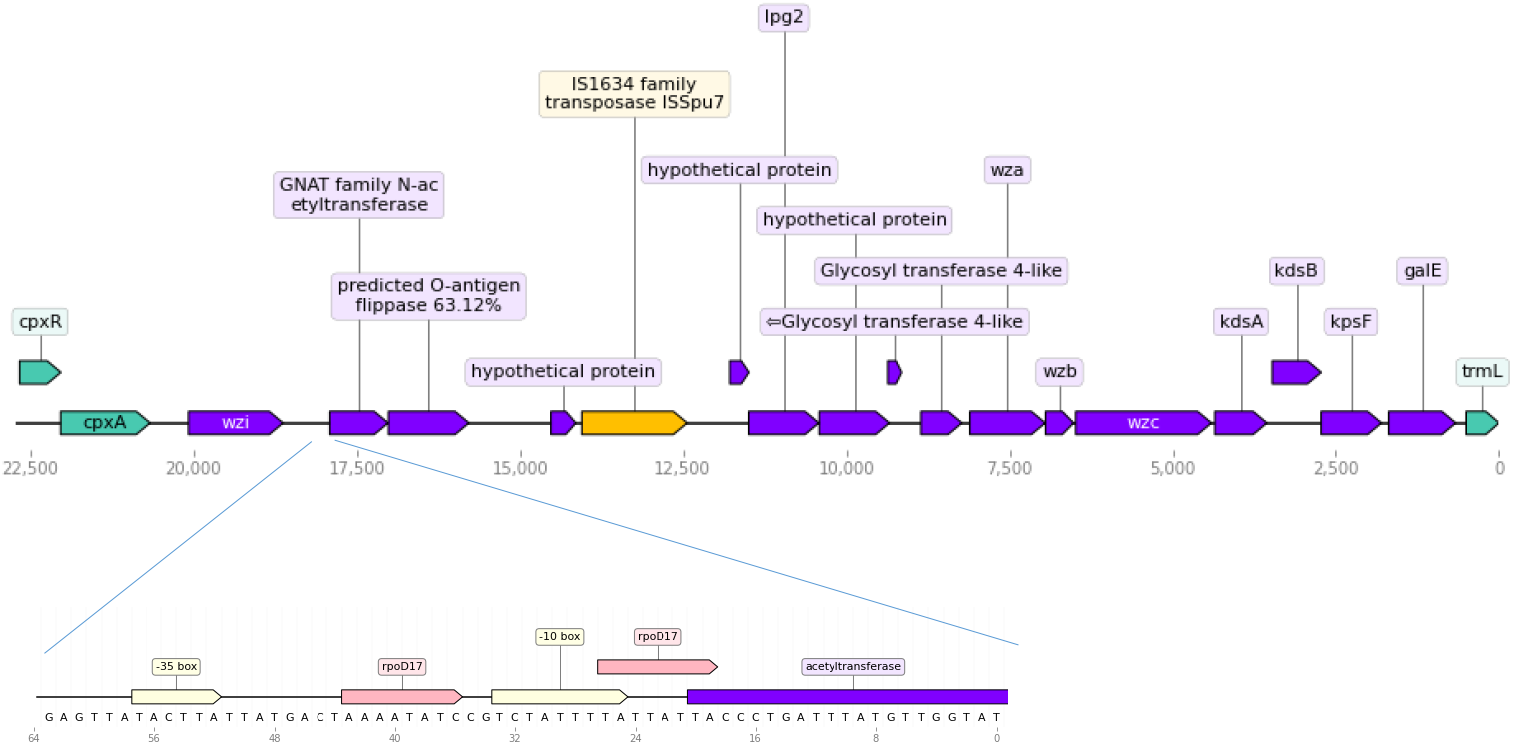
O-antigen composition of the *P. heimbachae* strain NCTC12003. The IS element is marked by yellow. Annotation of genes is based on Prokka.

We revealed several cases when insertion of an IS element into the O-antigen operon did not result in a gene disruption. Promoters were predicted in the upstream region of such IS elements. To check whether a candidate promoter was brought to operon together with an inserted IS element, we aligned the upstream and downstream regions of the IS element with the respective intergenic regions from operons of *Providencia* species with similar operon structure if any was found in our data (Fig. 8).

**Figure 8.**
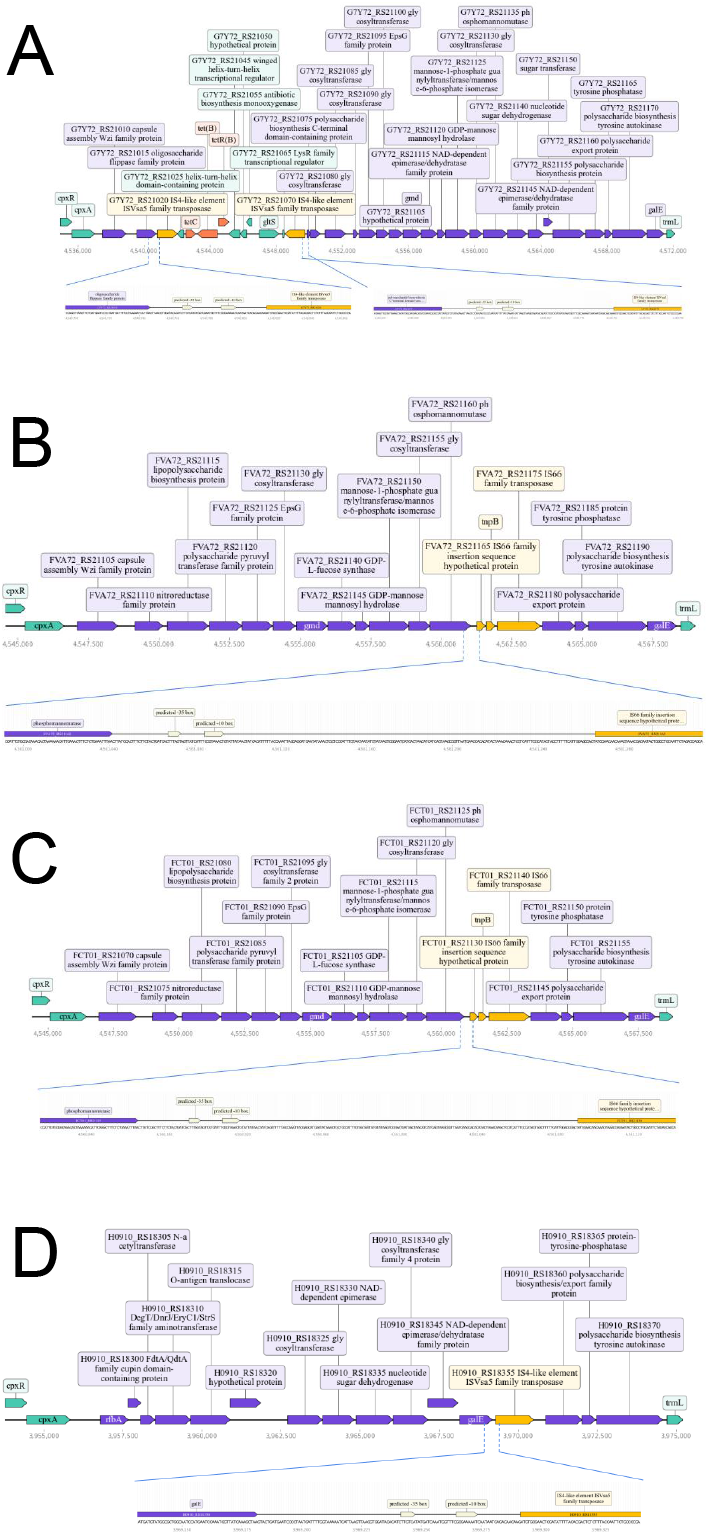
Organization of the O-antigen operon in *P. rettgeri* strain BML2496 (A), *Providencia sp*. 1709051003 (B), *P. rettgeri* strain Pr-15-2-50 (C), and *P. alcalifaciens* strain 1701003 (D). The IS element is marked yellow. Predicted promoter regions are coloured beige. For each gene annotated by PGAP, either gene name or locus tag and product are indicated.

We also were able to predict promoters within the upstream regions of IS elements causing gene disruptions in *P. rettgeri* strain PreM15758, *P. rustigianii* strain NCTC6933, *P. rustigianii* strain NCTC8113, and *P. heimbachae* strain NCTC12003 (Figure S1 in Supplementary materials). As with *P. rettgeri* strain BML2496, the promoter verification can not be provided because the composition of respective O-antigen operons was different from those in other strains without IS elements.

The structure of the O-antigen operon in *Providencia sp*. 1709051003 and *P. rettgeri* strain Pr-15-2-50 is similar to *Providencia sp*. 1701091, *Providencia sp*. 1701011, and *P. rettgeri* strain 2055 which have no IS elements in the respective operons. Comparing the intergenic regions of *Providencia sp*. 1709051003 and *P. rettgeri* strain Pr-15-2-50, we found that candidate promoter near the start of the first IS66 family transposase gene (Fig. 8 B, C) might correspond to conserved sites downstream of the phosphomutase gene. It could indicate that the IS element insertion was not associated with incorporating additional promoters into the operon.

Upstream region of the transposase from *P. alcalifaciens* 1701003 (Fig. 8 D) was compared with the intergenic region between *galE* and *wza* genes of *P. alcalifaciens* FDAARGOS 408. The predicted upstream promoter of the IS transposase was located within the inserted fragment, not in the common intergenic region (Figure S2 in Supplementary materials). We suggest that the hypothetical source of this candidate regulatory region is the IS element. These findings point towards the possible role of IS elements in changing gene regulation by bringing regulatory sequences such as promoters into the operon.

### Candidates for O-antigen genes

In this study, we have expanded the list of candidate O-antigen genes in *Providencia* (Table 2). In the O-antigen operon, we revealed in total twelve genes that have been described in *P. alcalifaciens* by Ovchinnikova [23] as a part of the O40 gene cluster. We detected homologs of *wpaD, wpaG*, and *wpaA* only in several *P. rettgeri* strains (Table S5 in Supplementary materials) and *Providencia sp*. 2.29. Gene annotated with Prokka as *mshA* coding for glycosyltransferase shared 67.85% similarity with *wpaB* of *P. alcalifaciens*. Another glycosyltransferase encoded by *tuaG* was similar to *wpaB*, exhibiting from about 66% up to 74% similarity depending on the organism. A homolog of *wza* gene was found in the O-antigen operon as an unknown gene which product function was predicted as polysaccharide biosynthesis/export family protein. Described gene was at 98.52% similar to *wza* in *P. alcalifaciens* strain FDAARGOS_408 and no less than at 71% in *P. rettgeri, P. stuartii*, and *P. heimbachae* strains. Remarkably, the gene *wza* was always colocalized with *wzb* and *wzc* genes forming a sustainable gene combination.

**Table 2:** Genes associated with O-antigen biosynthesis in *Providencia* species. Genes that have not been previously reported as O-antigen genes in *Providencia* species are marked with an asterisk.

| Gene name | Protein function |
| --- | --- |
| *galE* | UDP-glucose 4-epimerase |
| *rfbB* | dTDP-glucose 4,6-dehydratase |
| *rfbA* * | Glucose-1-phosphate thymidyltransferase |
| *fdtA* (*qdtA*) | Isomerase |
| *gmd* | GDP-mannose 4,6-dehydratase |
| *neuB* * | N,N'-diacetyllegionaminic acid synthase |
| *neuC* * | Polysialic acid biosynthesis protein |
| *arnC* | Glycosyltransferase |
| *gmm* | GDP-mannose mannosyl hydrolase |
| *rfbM* | Mannose-1-phosphate guanylyltransferase |
| *glmM* | Phosphoglucosamine mutase |
| *gnu* | Epimerase |
| *wbnH* | O-antigen biosynthesis glycosyltransferase |
| *wzx* | O-antigen flippase |
| *ugd* | UDP-glucose 6-dehydrogenase |
| *wbgU* * | UDP-N-acetylglucosamine 4-epimerase |
| *fdtB* | Aminotransferase |
| *fdtC* | Acetyltransferase |
| *wfaP* * | Glycosyltransferase |
| *tuaG* * | Glycosyltransferase |
| *wbbD* * | Galactosyltransferase |
| *rmlD* | Oxidoreductase |
| *rfbC* | Isomerase |
| *kdsA* | Transferase |

Table 2: Genes associated with O-antigen biosynthesis in *Providencia* species. Genes that have not been previously reported as O-antigen genes in *Providencia* species are marked with an asterisk. (Continued)
|  |  |
| --- | --- |
| *kdsB* | Nucleotidyltransferase |
| *kpsF* | Arabinose 5-phosphate isomerase |
| *wfgD* | Glycosyltransferase |
| *pglH* | Glycosyltransferase |
| *pglF* | Lyase |
| *arnB* | Aminotransferase |

*P. alcalifaciens* O40 gene cluster was bordered by the *cpxA* and *yibK (tmrL*) housekeeping genes according to the Ovchinnikova et al. study [23]. Due to the common boundaries, we can state that the operons we detected are indeed operons of O-antigens.

To the best of our knowledge, about a quarter of the identified genes within the O-antigen operons have not been previously reported as O-antigen genes in *Providencia* species (Table 2, genes indicated with an asterisk (* symbol)).

### Phylogenetic analysis

According to phylogenetic analysis, each *Providencia* species formed a distinct cluster on the phylogenetic tree. Species-level classification of unknown *Providencia* can be predicted based on the *galE* phylogeny tree. We assume that *Providencia sp*. 1701091, *Providencia sp*. 1701011, and *Providencia sp*. 1709051003 belong to the *P. rettgeri*, and *Providencia sp*. 229 could correspond to *P. stuartii* species (Fig. 9, Table S4 in Supplementary materials).

**Figure 9.**
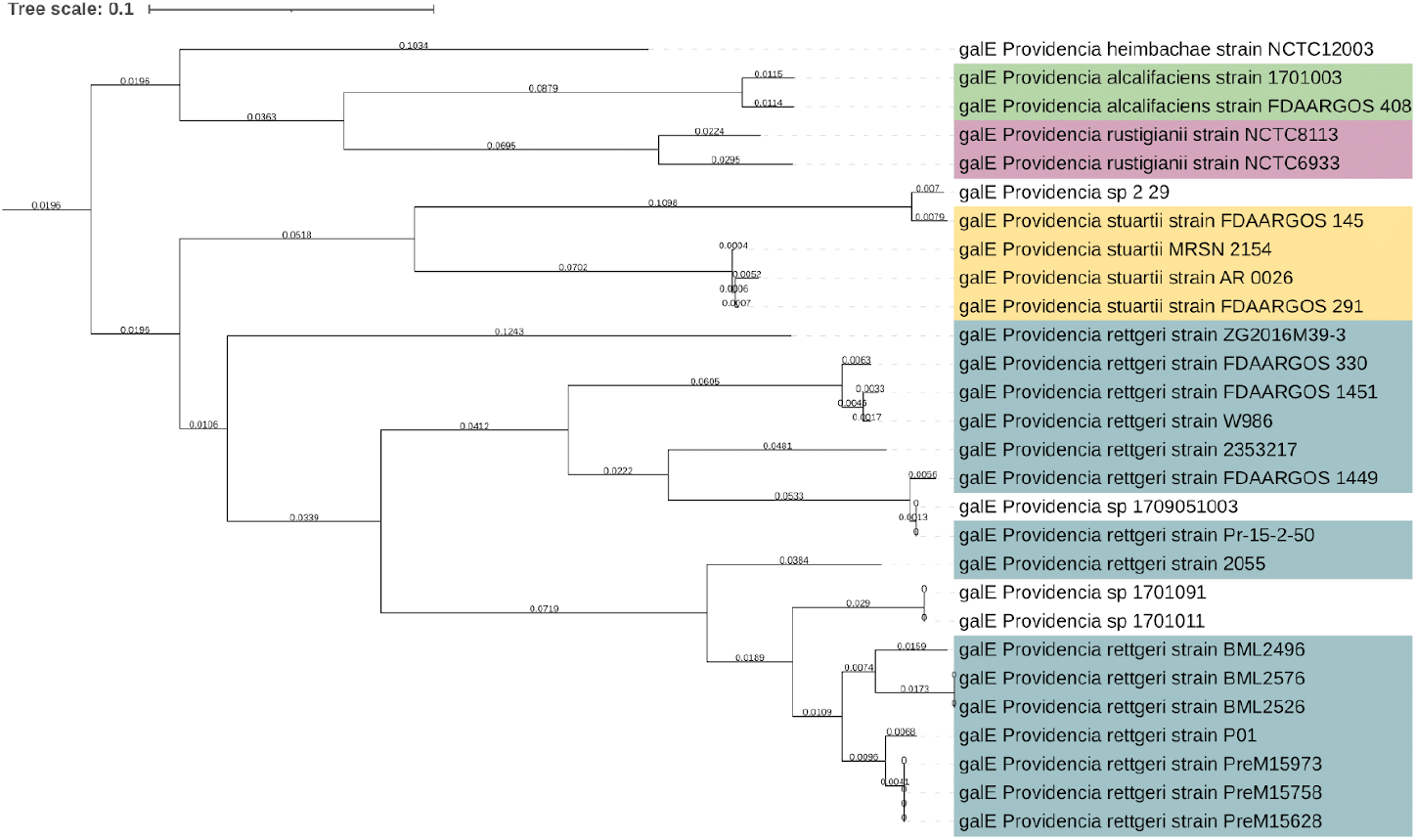
Phylogenetic tree of *galE* gene. Labels are colored by *Providencia* species. Unknown *Providencia* species are marked white.

Clustering of species on phylogenetic *galE* gene trees predominantly correlated with differences in the structure of O-antigens operons flanked by *cpxR* and *trml* genes. For instance, by the composition of the operon, *P. rettgeri* strain ZG2016M39-3 was most similar to *P. alcalifaciens* but not to *P. rettgeri* (Fig. 9, Fig. 10). A distinctive feature of this operon in *P. alcalifaciens, P. rustigianii* strains, and *P. rettgeri* strain ZG2016M39-3 was the absence of the *wzi* gene. On the phylogenetic tree, the respective species of *Providencia* were grouped together and formed a separate clade (Fig. 9).

**Figure 10.**
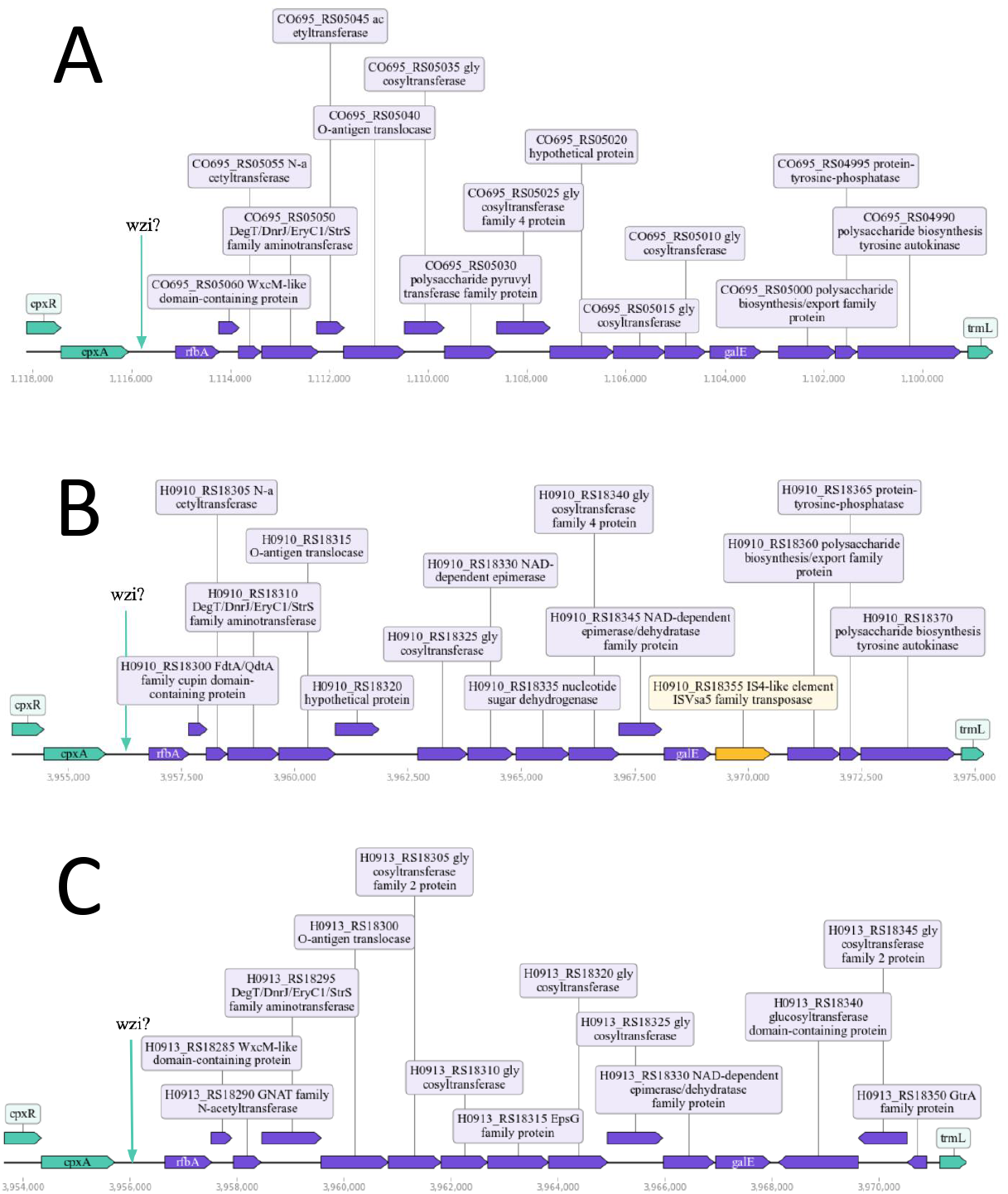
Organization of the O-antigen operon in *P. alcalifaciens* strain FDAAR-GOS_408 (A), *P. alcalifaciens* strain 1701003 (B), and *P. rettgeri* strain ZG2016M39-3 (C). The *wzi* gene is absent between the *cpxRA* and O-antigen operons. For each gene annotated by PGAP, either gene name or locus tag and product are indicated.

## Discussion

The provided study aimed to clarify the O-antigen operons composition for *Providencia* species. While study performing, the unique pipeline enabled to reveal O-antigen operons in *Providencia* species was created. Compared to existing ones, it relies on the operon level and has several stages of results verification. The first step checks the annotation output, while the second verifies positions of the operon boundaries. Only a combination of features provides reliable information about the localization and composition of the operon in the genome. Subsequently this pipeline can be applied to search for any operon structures.

The first pipeline step allows to minimize the annotation mistakes. Prokka identifies all the main products, while PGAP provides control of Prokka results. EggNOG and local alignment add information about unannotated products from the previous step. In this way, more accurate data is obtained. Since Operon mapper defines operon boundaries on only intergenic distance and functional relationships, a check of operons’ boundary was included. Two tools were added to verificate promoter and terminator positions, which mark the start and the end of the operon. This necessity became clear after the comparison analysis provided by Assaf et al [44].

The pipeline identified new features in the O-antigen composition of *Providencia* species. Initially somatic antigen was described by Ovchinnikova as a set of genes located between *cpxA* and *yibK* genes. The idea of its composition has not been changed due to the gene level of analysis [16, 45]. Switching to the operonic level of search discovers new genes connected with the O-antigen biosynthesis [46]. Another finding is the O-antigen operon with a transposon insertion. By it, *P. rettgeri* strain BML2496 may have become resistant to tetracycline. The presence of IS elements in the O-antigen has been shown for five *Providencia* strains belonging to four species.

Current work may shed light on some taxonomic misunderstandings. The phylogenetic tree based on O-antigen operons led to several clarifications in the *Providencia* taxonomy. Most of the unknown species are grouped together with the *P. rettgeri* strain, while one of them is clustered with the *P. stuartii* branch. The *P. rettgeri* strain ZG2016M39-3 is closer to *P. rustigianii* or *P. alcalifaciens* rather than other *P. rettgeri* bacteria. Classification mistakes are possible in this case. Coming to the end, there are still many mysteries in the *Providencia* O-antigen composition. To the results obtained, there is a quite reliable link between the lifestyle and O-antigen composition for *Providencia* species. We plan to test the same hypothesis on other bacterial species with contrasting life types to strengthen or refute this idea.

## Supplementary materials

All supporting information can be found here:

Supplementary Table S1
Supplementary Table S2
Supplementary Table S3
Supplementary Table S4
Supplementary Table S5
Supplementary Figure S1
Supplementary Figure S2

## Funding

Aleksey Komissarov was financially supported by the ITMO Fellowship and Professorship Program.

## Notes

### Competing Interest Statement

The authors have declared no competing interest.

