## Supplementary material for "Identification and comparison of somatic antigen composition for bacteria from *Providencia* genus": ./supplement/figureS1.pdf

**Figure S1.** Organization of the O-antigen operon in *P. rettgeri* strain PreM15758 (A), *P. stuartii* strain AR_0026 (B), *P. rustigianii* strain NCTC6933 (C), *P. rustigianii* strain NCTC8113 (D), and *P. heimbachae* strain NCTC12003 (E). The IS element is marked yellow. Predicted promoter regions are colored beige. For each gene annotated by PGAP, locus tag and product are indicated.

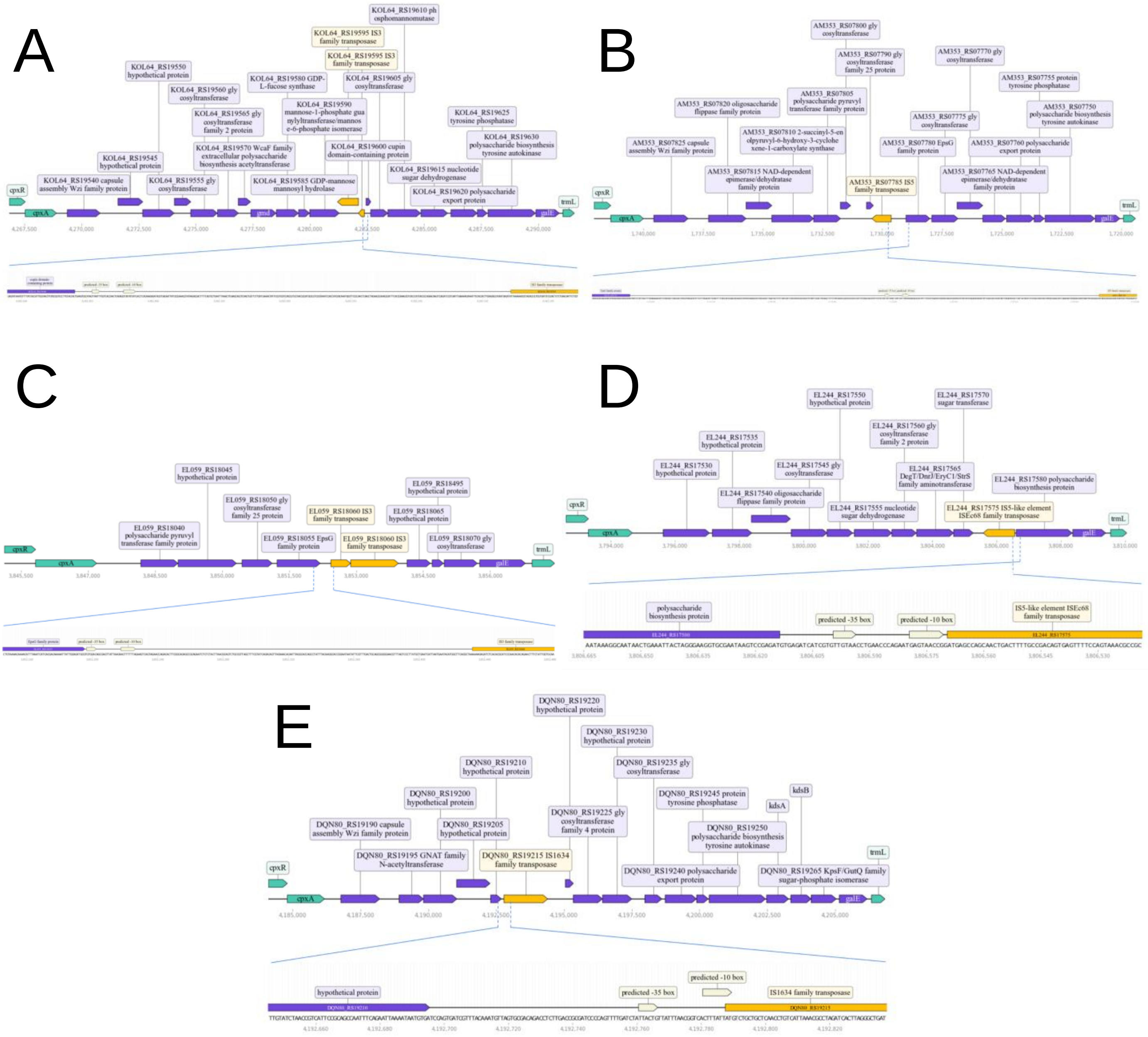
