## Supplementary material for "Identification and comparison of somatic antigen composition for bacteria from *Providencia* genus": ./supplement/figureS2.pdf

**Figure S2.** Sequence comparison between intergenic region of *galE*/*wza* genes of *P. alcalifaciens* strain FDAARGOS_408 (upper sequence) and upstream region of transposase from *P. alcalifaciens* strain 1701003 (lower sequence). Alignment was done by T-COFFEE (version 11) with default parameters. Stop codon is marked red, start codon is marked green, predicted promoter is marked beige.

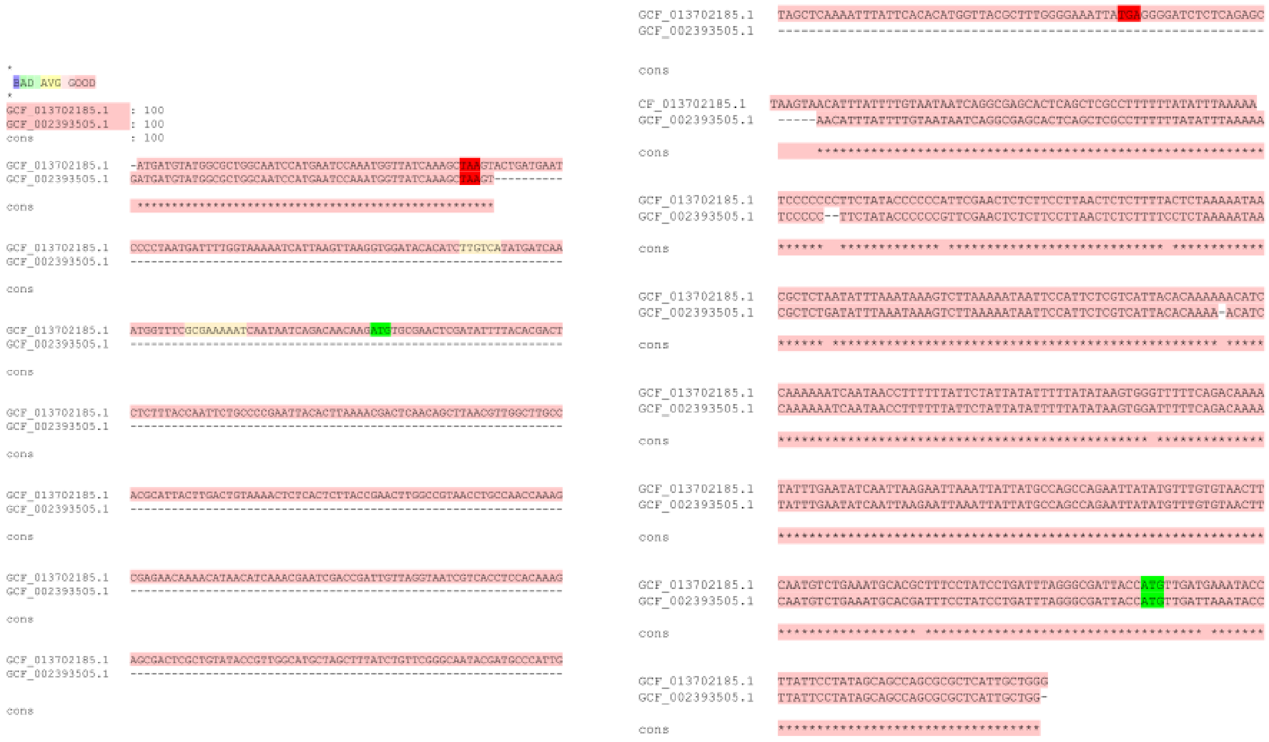
