## Supplementary material for "Identification and comparison of somatic antigen composition for bacteria from *Providencia* genus": ./supplement/tableS1.pdf

**Table S1**. Summary of assemblies

| RefSeq Assembly Accession | Organism Name | Total sequence length | # contigs | N50 | L50 |
|---|---|---|---|---|---|
| GCF_000259175.1 | *Providencia stuartii* MRSN 2154 | 4402109 | 1 | 4402109 | 1 |
| GCF_001558855.2 | *Providencia stuartii* strain FDAARGOS_145 | 4305109 | 1 | 4305109 | 1 |
| GCF_002393505.1 | *Providencia alcalifaciens* strain FDAARGOS_408 | 3990106 | 1 | 3990106 | 1 |
| GCF_002947315.1 | *Providencia stuartii* strain AR_0026 | 4368167 | 1 | 4368167 | 1 |
| GCF_002983665.1 | *Providencia stuartii* strain FDAARGOS_291 | 4394489 | 1 | 4394489 | 1 |
| GCF_002984195.1 | *Providencia rettgeri* strain FDAARGOS_330 | 4518601 | 1 | 4518601 | 1 |
| GCF_005155965.1 | *Providencia rettgeri* strain Pr-15-2-50 | 4649193 | 1 | 4649193 | 1 |
| GCF_010318885.1 | *Providencia rettgeri* strain BML2496 | 4651003 | 1 | 4651003 | 1 |
| GCF_010319105.1 | *Providencia rettgeri* strain BML2526 | 4342905 | 1 | 4342905 | 1 |
| GCF_010319405.1 | *Providencia rettgeri* strain BML2576 | 4351300 | 1 | 4351300 | 1 |
| GCF_011045615.1 | *Providencia* sp. 1701011 | 4596310 | 1 | 4596310 | 1 |
| GCF_011045635.1 | *Providencia* sp. 1701091 | 4623927 | 1 | 4623927 | 1 |
| GCF_011045655.1 | *Providencia* sp. 1709051003 | 4649235 | 1 | 4649235 | 1 |
| GCF_013694385.1 | *Providencia rettgeri* strain P01 | 4601020 | 1 | 4601020 | 1 |
| GCF_013702025.1 | *Providencia rettgeri* strain 2055 | 4674814 | 1 | 4674814 | 1 |

**Table S1**. Summary of assemblies

| | | | | | |
|---|---|---|---|---|---|
| GCF_013702185.1 | *Providencia alcalifaciens* strain 1701003 | 4033976 | 1 | 4033976 | 1 |
| GCF_013702245.1 | *Providencia rettgeri* strain 2353217 | 4337992 | 1 | 4337992 | 1 |
| GCF_013702265.1 | *Providencia rettgeri* strain ZG2016M39-3 | 4023756 | 1 | 4023756 | 1 |
| GCF_015832305.1 | *Providencia* sp. 2.29 | 4547792 | 1 | 4547792 | 1 |
| GCF_018771265.1 | *Providencia rettgeri* strain W986 | 4515465 | 1 | 4515465 | 1 |
| GCF_018861215.1 | *Providencia rettgeri* strain PreM15973 | 4367913 | 1 | 4367913 | 1 |
| GCF_018861235.1 | *Providencia rettgeri* strain PreM15628 | 4368530 | 1 | 4368530 | 1 |
| GCF_018861255.1 | *Providencia rettgeri* strain PreM15758 | 4368564 | 1 | 4368564 | 1 |
| GCF_019047885.1 | *Providencia rettgeri* strain FDAARGOS 1449 | 4777252 | 1 | 4777252 | 1 |
| GCF_019048545.1 | *Providencia rettgeri* strain FDAARGOS 1451 | 4383816 | 1 | 4383816 | 1 |
| GCF_900475855.1 | *Providencia heimbachae* strain NCTC12003 | 4286000 | 1 | 4286000 | 1 |
| GCF_900635875.1 | *Providencia rustigianii* strain NCTC6933 | 3913850 | 1 | 3913850 | 1 |
| GCF_900637755.1 | *Providencia rustigianii* strain NCTC8113 | 3864394 | 1 | 3864394 | 1 |
