## Supplementary material for "Identification and comparison of somatic antigen composition for bacteria from *Providencia* genus": ./supplement/tableS2.pdf

**Table S2**. The length of the O-antigen operon in all *Providencia* species used in the study

| **RefSeq Assembly Accession** | **Organism name** | **O-antigen operon length (*cpxA-trmL*)** |
|---|---|---|
| GCF_000259175.1 | *Providencia stuartii* strain MRSN 2154 | 19063 |
| GCF_002983665.1 | *Providencia stuartii* strain FDAARGOS_291 | 19435 |
| GCF_002947315.1 | *Providencia stuartii* strain AR_0026 | 22649 |
| GCF_001558855.2 | *Providencia stuartii* strain FDAARGOS_145 | 22931 |
| GCF_015832305.1 | *Providencia sp.* 2.29 | 22913 |
| GCF_002393505.1 | *Providencia alcalifaciens* strain FDAARGOS_408 | 19550 |
| GCF_013702185.1 | *Providencia alcalifaciens* strain 1701003 | 21446 |
| GCF_013702265.1 | *Providencia rettgeri(alcalifaciens?)* strain ZG2016M39-3 | 18006 |
| GCF_900635875.1 | *Providencia rustigianii* strain NCTC6933 | 12312 |
| GCF_900637755.1 | *Providencia rustigianii* strain NCTC8113 | 17440 |
| GCF_900475855.1 | *Providencia heimbachae* strain NCTC12003 | 22732 |
| GCF_010319105.1 | *Providencia rettgeri* strain BML2526 | 19278 |
| GCF_010319405.1 | *Providencia rettgeri* strain BML2576 | 19278 |
| GCF_002984195.1 | *Providencia rettgeri* strain FDAARGOS_330 | 20253 |
| GCF_018771265.1 | *Providencia rettgeri* strain W986 | 20487 |
| GCF_019048545.1 | *Providencia rettgeri* strain FDAARGOS 1451 | 20488 |
| GCF_013702245.1 | *Providencia rettgeri* strain 2353217 | 23460 |
| GCF_018861215.1 | *Providencia rettgeri* strain PreM15973 | 23477 |
| GCF_018861235.1 | *Providencia rettgeri* strain PreM15628 | 23477 |
| GCF_013694385.1 | *Providencia rettgeri* strain P01 | 23624 |
| GCF_005155965.1 | *Providencia rettgeri* strain Pr-15-2-50 | 24421 |
| GCF_018861255.1 | *Providencia rettgeri* strain PreM15758 | 24742 |

**Table S2**. The length of the O-antigen operon in all *Providencia* species used in the study

| | | |
|---|---|---|
| GCF_019047885.1 | *Providencia rettgeri* strain FDAARGOS 1449 | 26642 |
| GCF_013702025.1 | *Providencia rettgeri* strain 2055 | 26781 |
| GCF_010318885.1 | *Providencia rettgeri* strain BML2496 | 37156 |
| GCF_011045615.1 | *Providencia sp.* 1701011 | 28967 |
| GCF_011045635.1 | *Providencia sp.* 1701091 | 28967 |
| GCF_011045655.1 | *Providencia sp.* 1709051003 | 24421 |
