## Supplementary material for "Identification and comparison of somatic antigen composition for bacteria from *Providencia* genus": ./supplement/tableS3.pdf

**Table S4.1**. Composition of O-antigen operons based on Prokka annotation

| **Organism name** | **Operon composition** |
|---|---|
| *Providencia stuartii* MRSN 2154 | 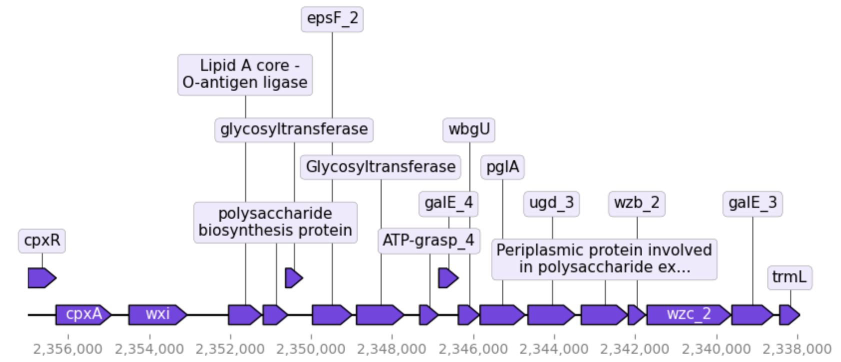 |
| *Providencia stuartii* FDAARGOS_145 | 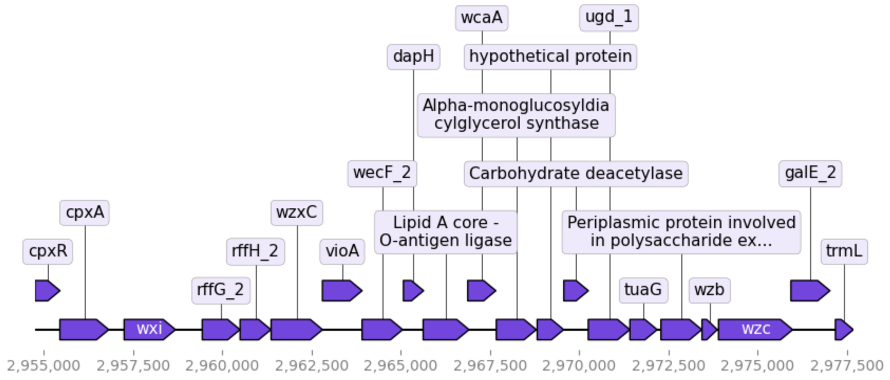 |
| *Providencia alcalifaciens* FDAARGOS_408 | 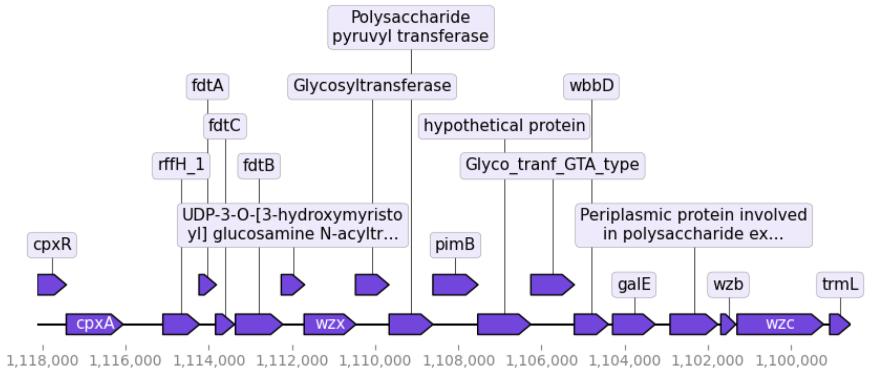 |

**Table S4.1**. Composition of O-antigen operons based on Prokka annotation

<table>
<tr><td>Providencia stuartii AR_0026</td><td>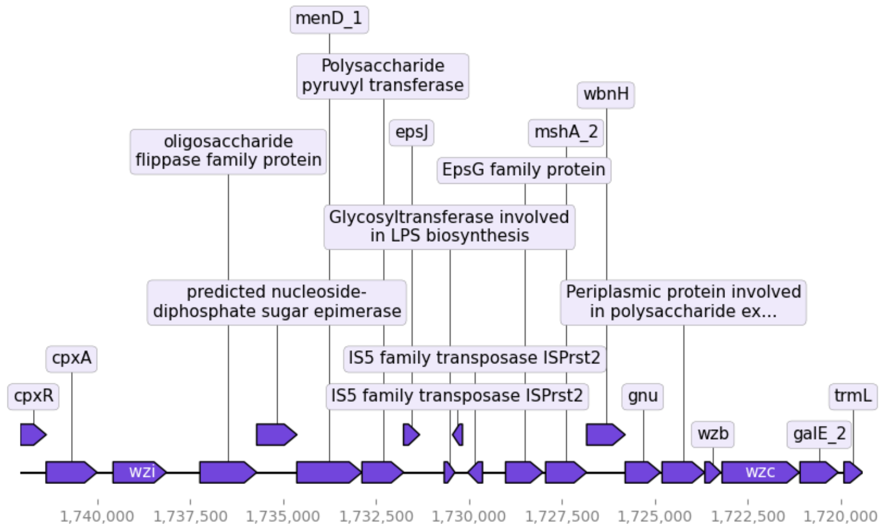
</td></tr>
<tr><td>Providencia stuartii FDAARGOS_291</td><td>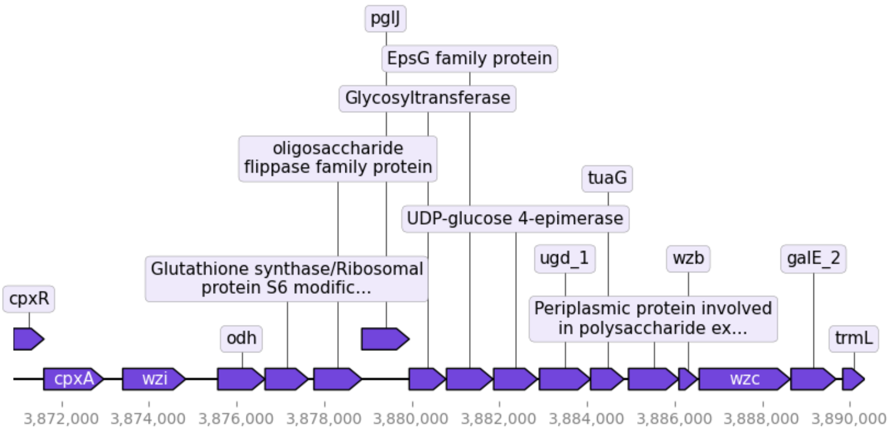
</td></tr>
<tr><td>Providencia rettgeri FDAARGOS_330</td><td>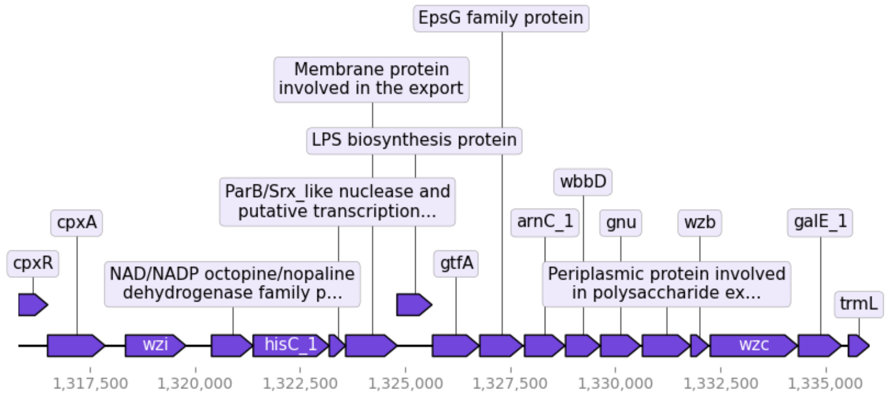
</td></tr>
</table>

**Table S4.1**. Composition of O-antigen operons based on Prokka annotation

| | |
|---|---|
| *Providencia rettgeri* Pr-15-2-50 | 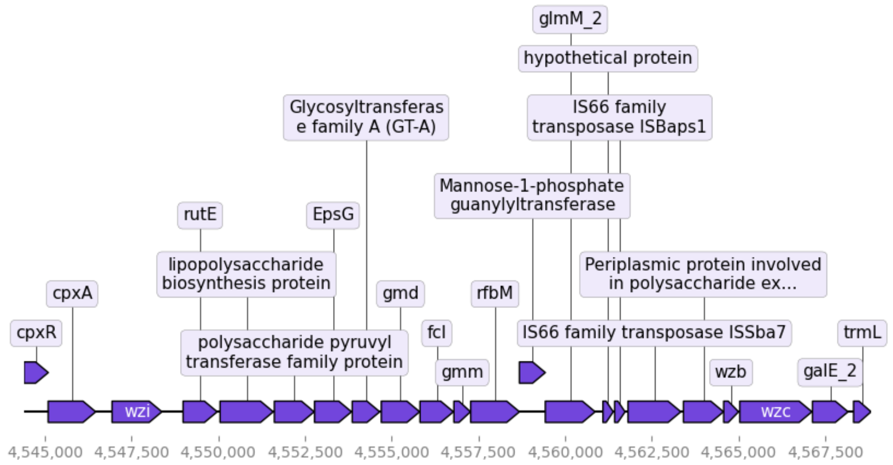 |
| *Providencia rettgeri* BML2496 | 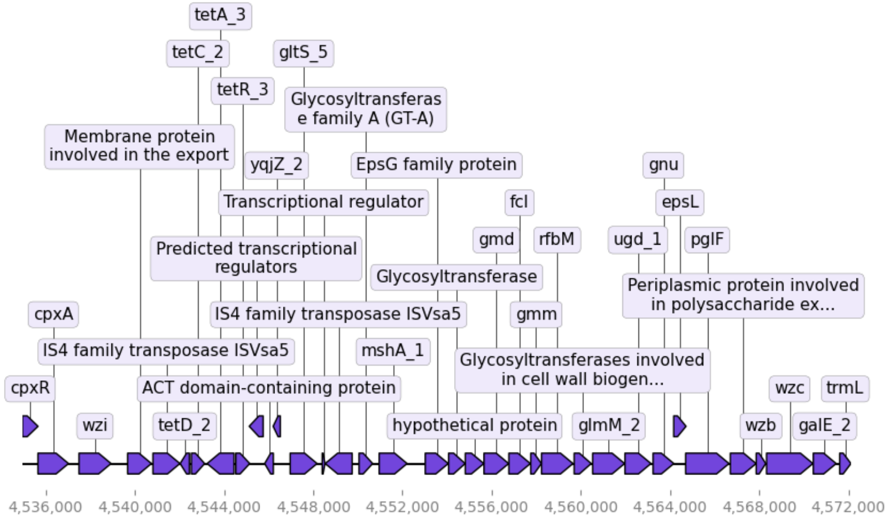 |
| *Providencia rettgeri* BML2526 | 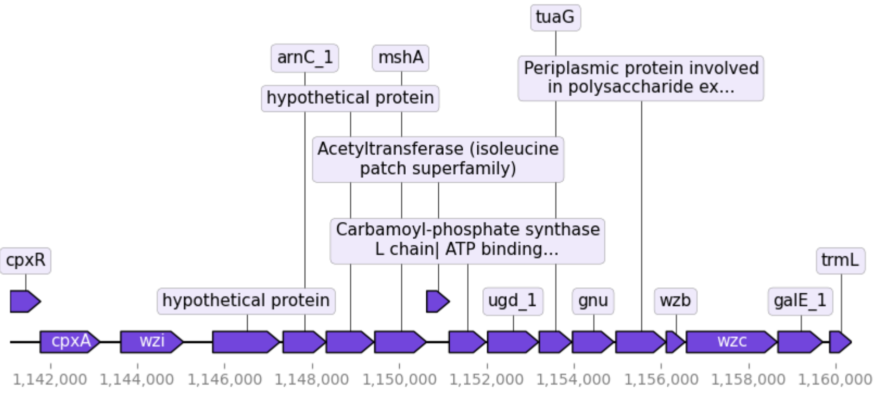 |

**Table S4.1**. Composition of O-antigen operons based on Prokka annotation

<table>
<tr><td>*Providencia rettgeri* BML2576</td><td>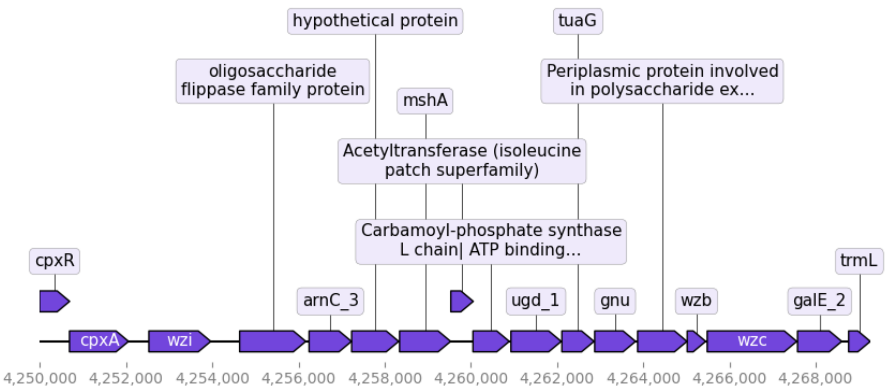
</td></tr>
<tr><td>*Providencia sp.* 1701011</td><td>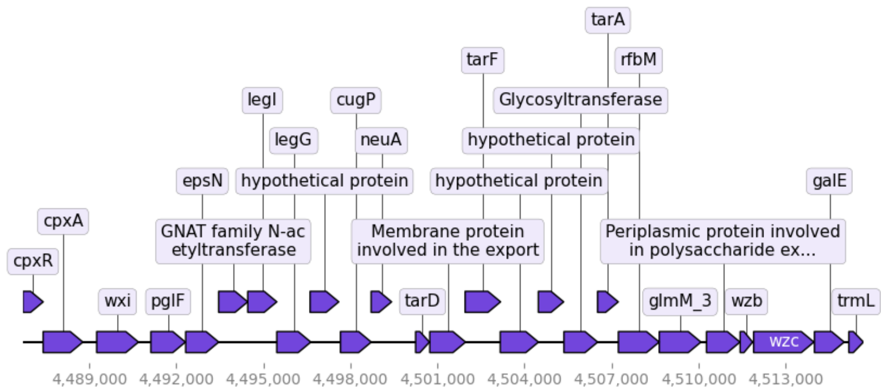
</td></tr>
<tr><td>*Providencia sp.* 1701091</td><td>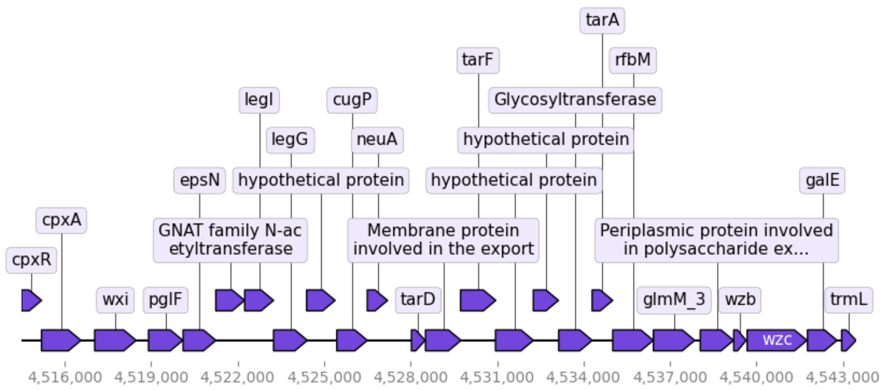
</td></tr>
</table>

**Table S4.1**. Composition of O-antigen operons based on Prokka annotation

<table>
<tr><td>Providencia sp.<br>1709051003</td><td>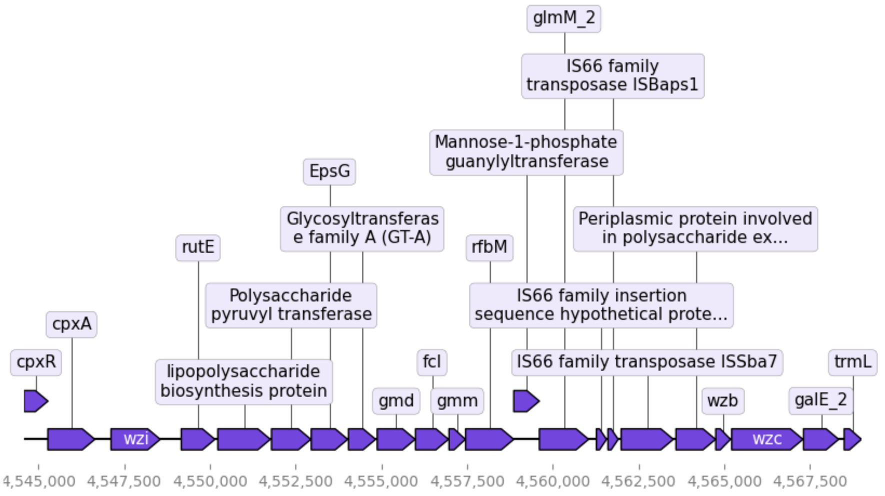
</td></tr>
<tr><td>Providencia rettgeri<br>P01</td><td>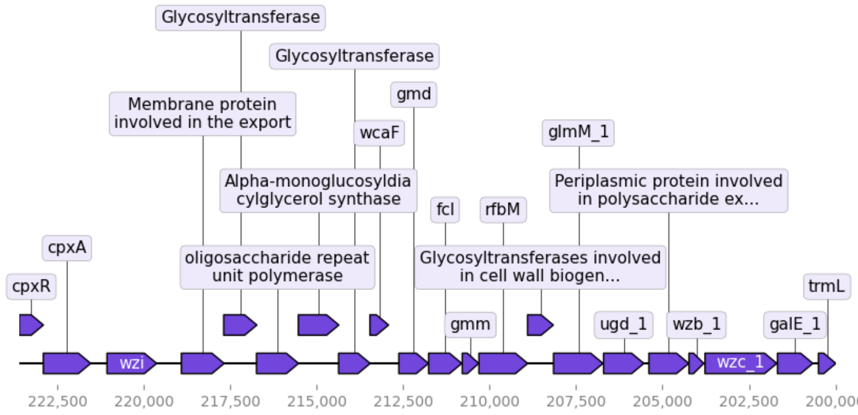
</td></tr>
<tr><td>Providencia rettgeri<br>2055</td><td>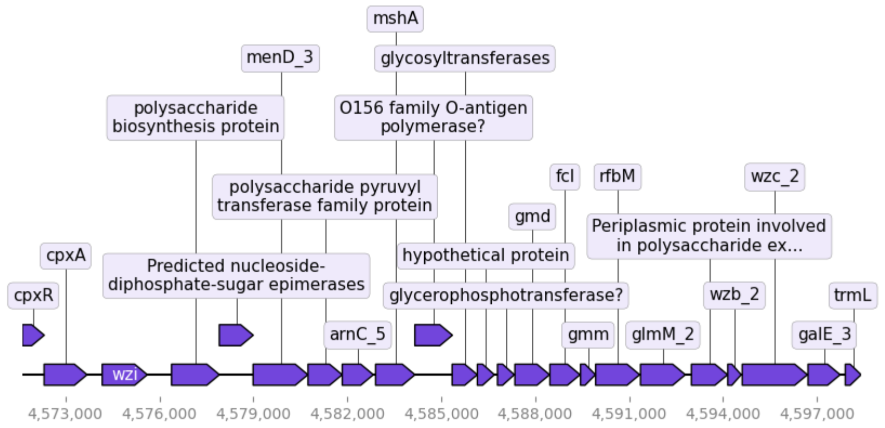
</td></tr>
</table>

**Table S4.1**. Composition of O-antigen operons based on Prokka annotation

| | |
|---|---|
| *Providencia alcalifaciens* 1701003 | 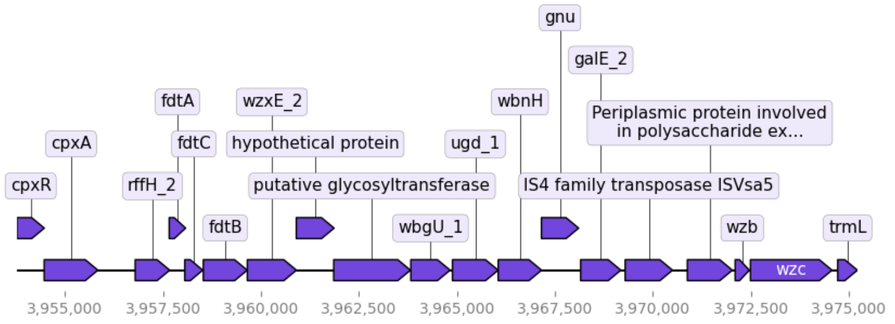 |
| *Providencia rettgeri* 2353217 | 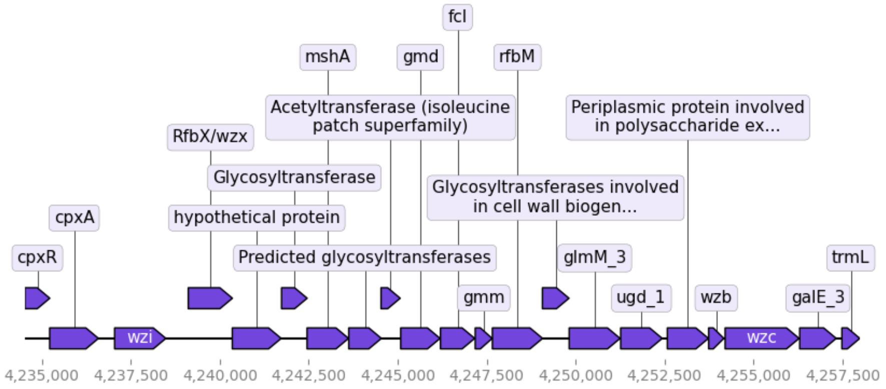 |
| *Providencia rettgeri* ZG2016M39-3 | 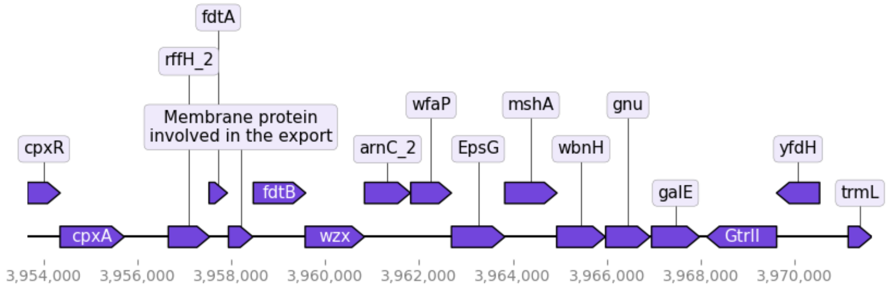 |
| *Providencia sp.* 2.29 | 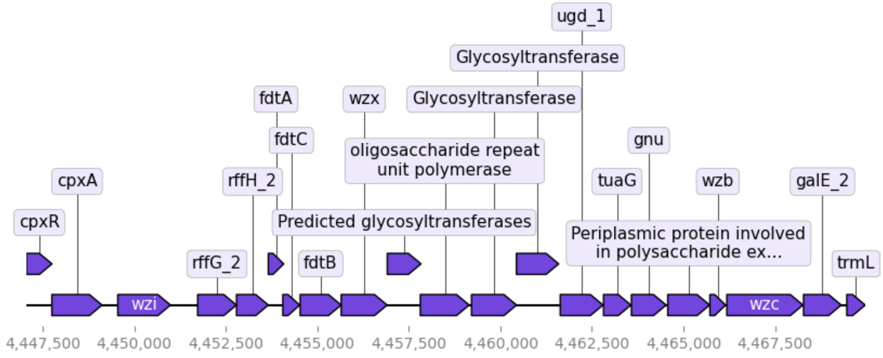 |

**Table S4.1**. Composition of O-antigen operons based on Prokka annotation

| | |
|---|---|
| *Providencia rettgeri* W986 | 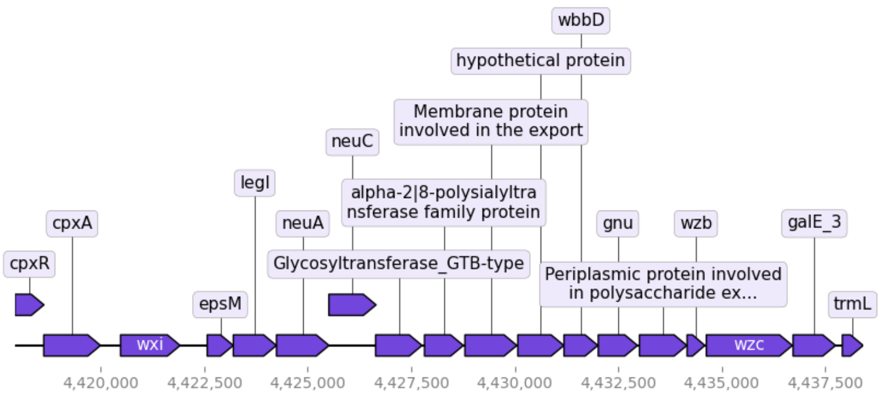 |
| *Providencia rettgeri* PreM15973 | 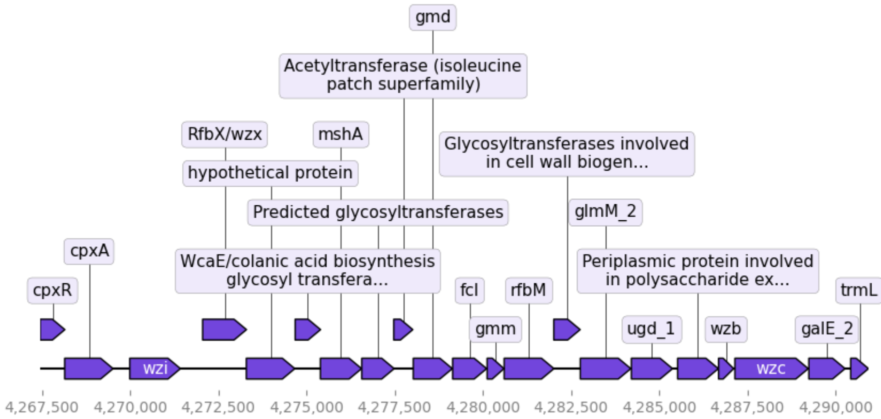 |
| *Providencia rettgeri* PreM15628 | 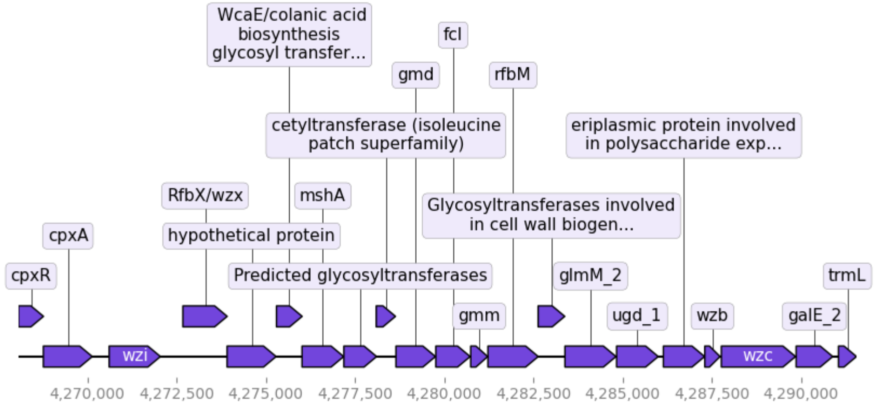 |
| *Providencia rettgeri* PreM15758 | 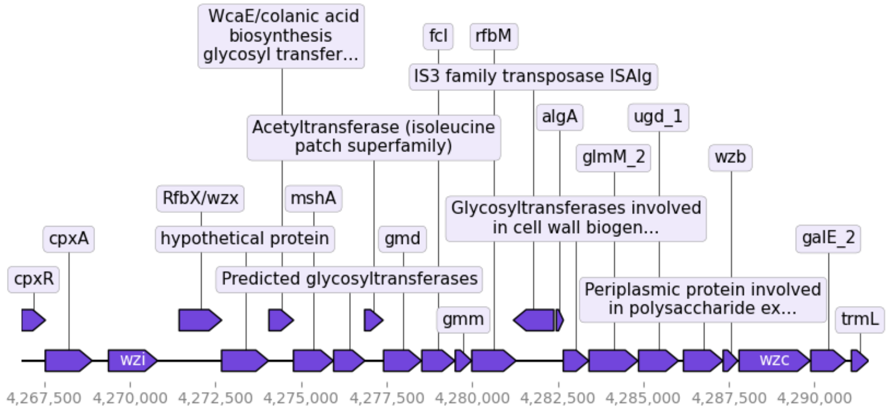 |

**Table S4.1**. Composition of O-antigen operons based on Prokka annotation

<table>
<tr><td>

*Providencia rettgeri* FDAARGOS 1449

</td><td>

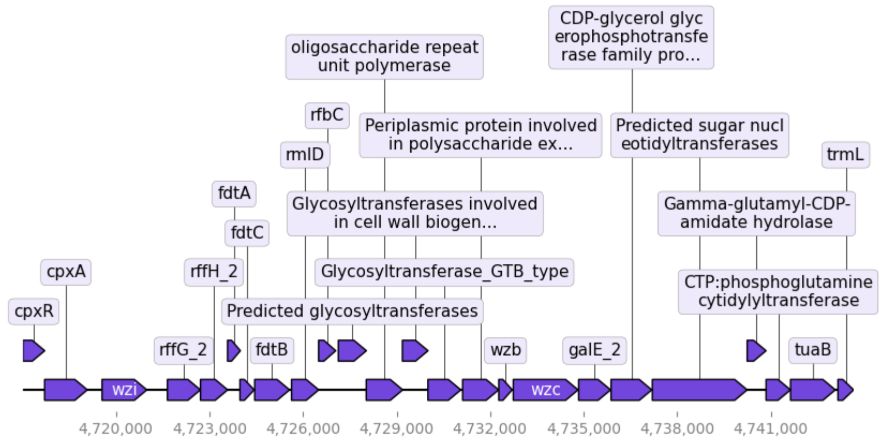


</td></tr>
<tr><td>

*Providencia rettgeri* FDAARGOS 1451

</td><td>

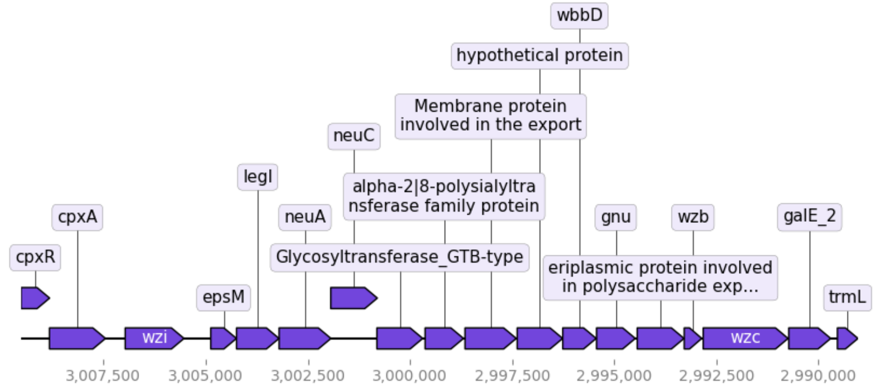


</td></tr>
<tr><td>

*Providencia heimbachae* NCTC12003

</td><td>

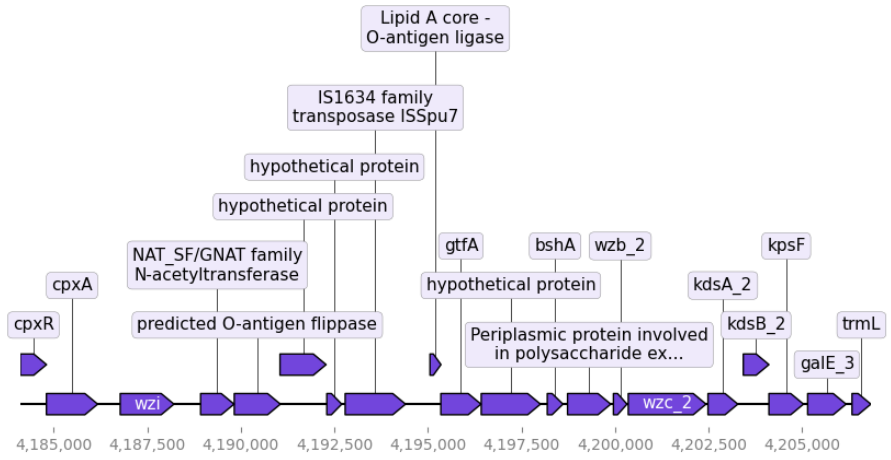


</td></tr>
<tr><td>

*Providencia rustigianii* NCTC6933

</td><td>

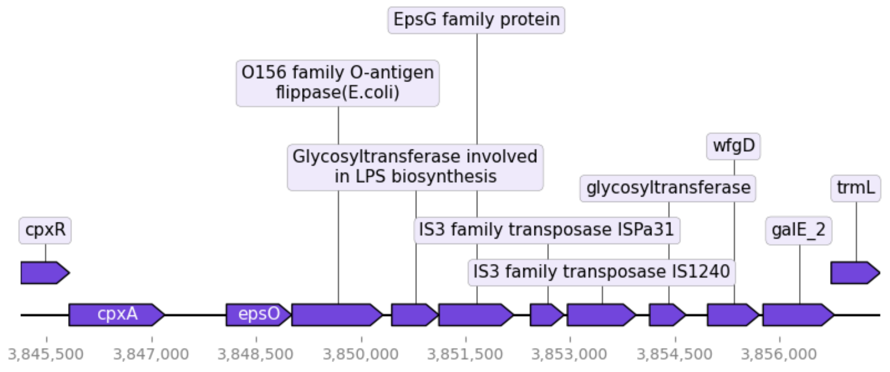


</td></tr>
</table>

**Table S4.1**. Composition of O-antigen operons based on Prokka annotation

<table>
<tr><td>Providencia rustigianii<br>NCTC8113</td><td>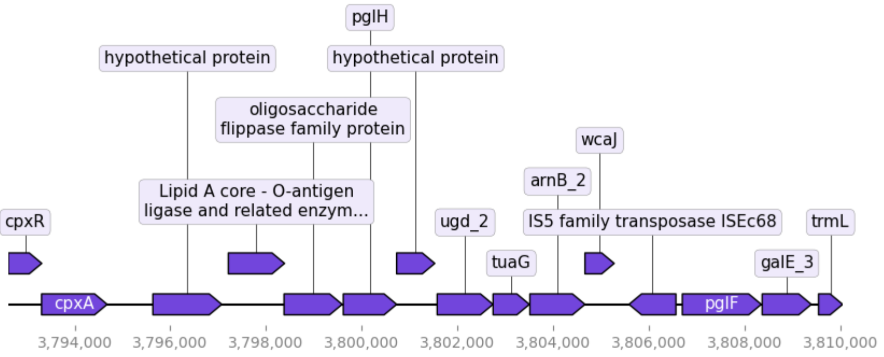
</td></tr>
</table>

**Table S4.2**. Composition of O-antigen operons based on PGAP annotation

<table>
<tr><th>Organism Name</th><th>Operon composition</th></tr>
<tr><td>Providencia stuartii<br>MRSN 2154</td><td>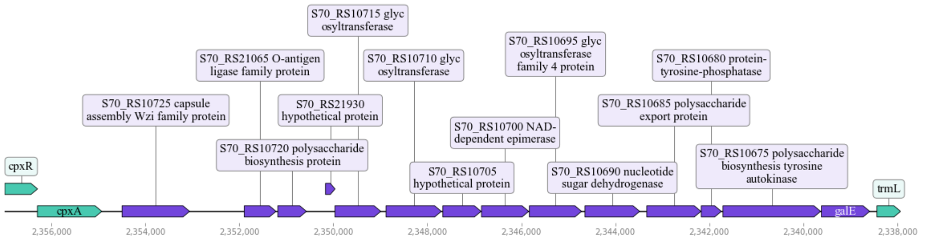
</td></tr>
<tr><td>Providencia stuartii<br>FDAARGOS_145</td><td>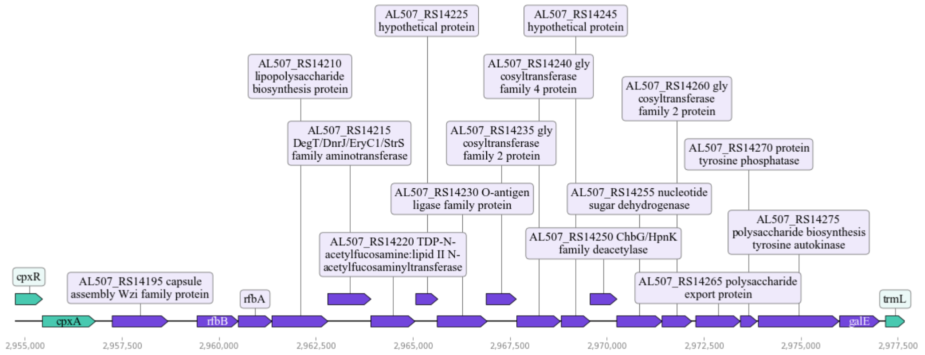
</td></tr>
<tr><td>Providencia alcalifaciens strain<br>FDAARGOS_408</td><td>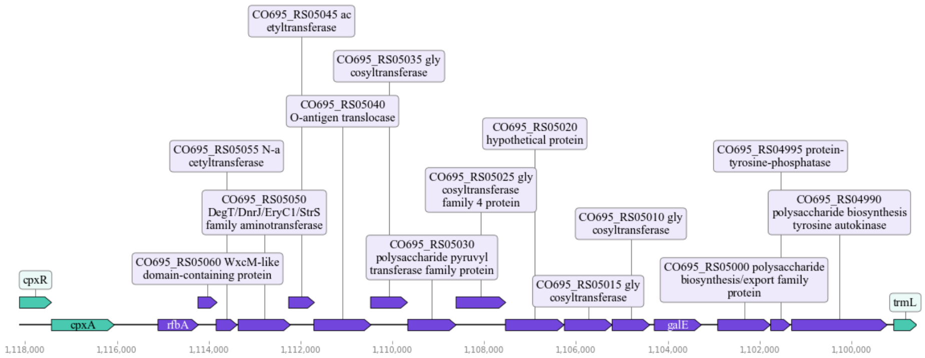
</td></tr>
</table>

**Table S4.2**. Composition of O-antigen operons based on PGAP annotation

<table>
<tr><td>Providencia stuartii AR_0026</td><td>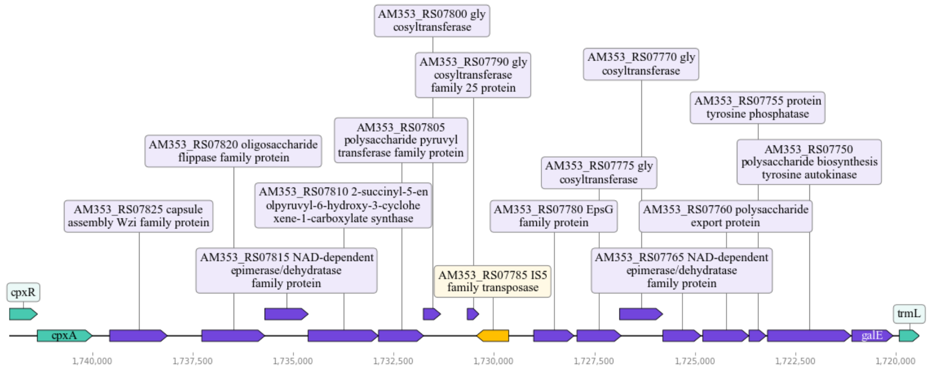
</td></tr>
<tr><td>Providencia stuartii FDAARGOS_291</td><td>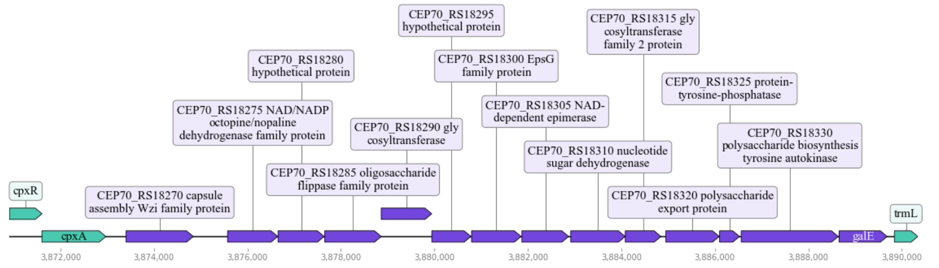
</td></tr>
<tr><td>Providencia rettgeri FDAARGOS_330</td><td>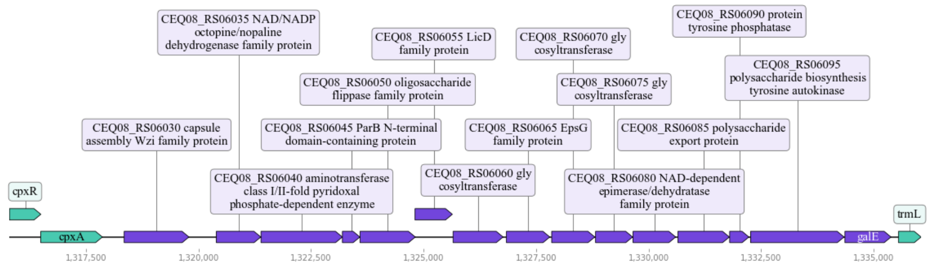
</td></tr>
<tr><td>Providencia rettgeri Pr-15-2-50</td><td>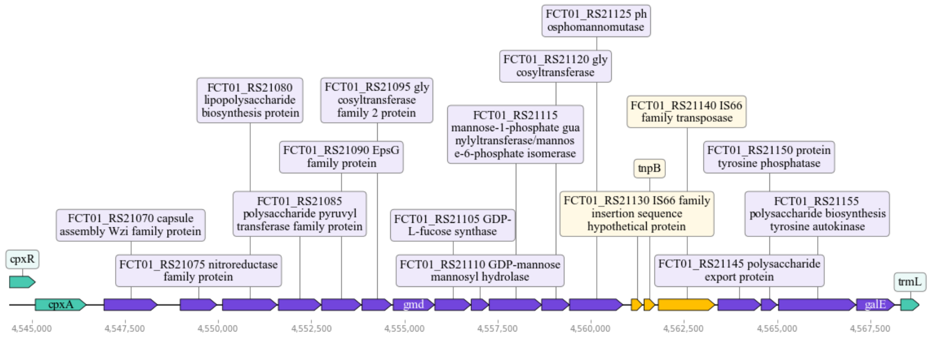
</td></tr>
</table>

**Table S4.2**. Composition of O-antigen operons based on PGAP annotation

| | |
|---|---|
| *Providencia rettgeri* BML2496 | 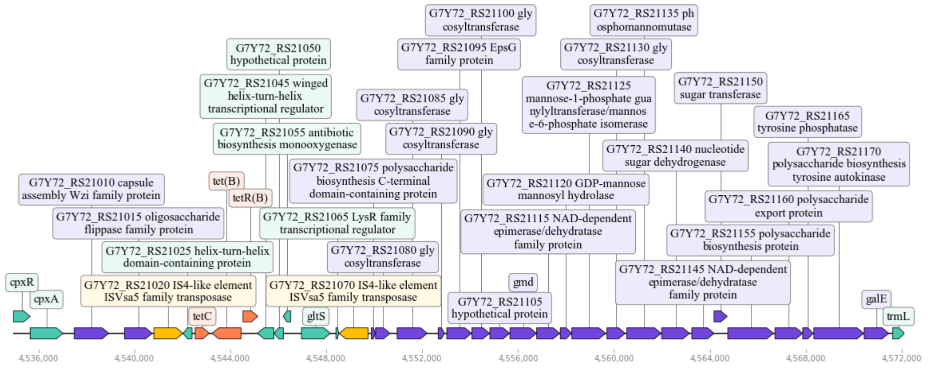 |
| *Providencia rettgeri* BML2526 | 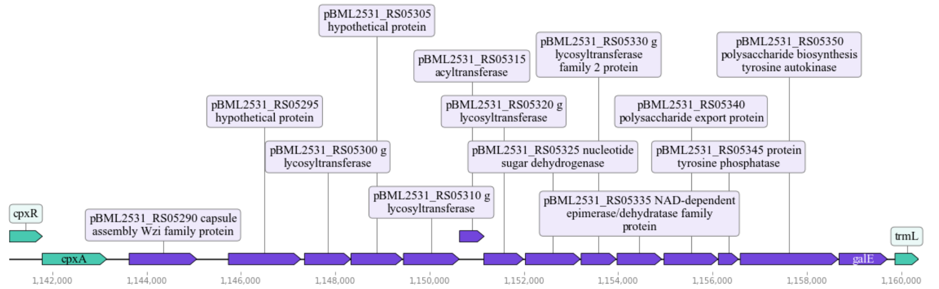 |
| *Providencia rettgeri* BML2576 | 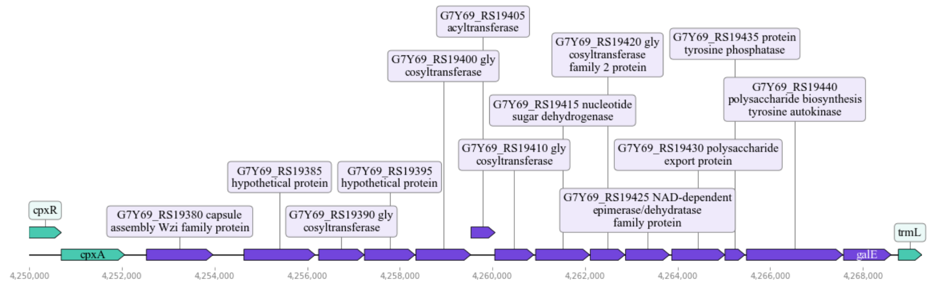 |
| *Providencia* sp. 1701011 | 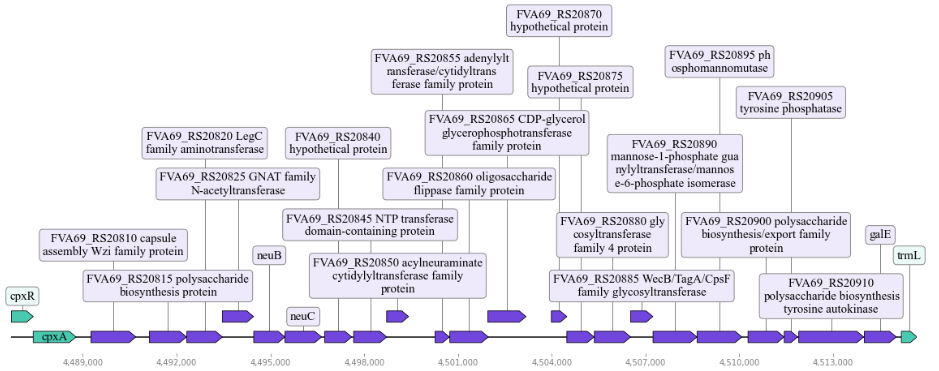 |

**Table S4.2**. Composition of O-antigen operons based on PGAP annotation

<table>
<tr><td>Providencia sp. 1701091</td><td>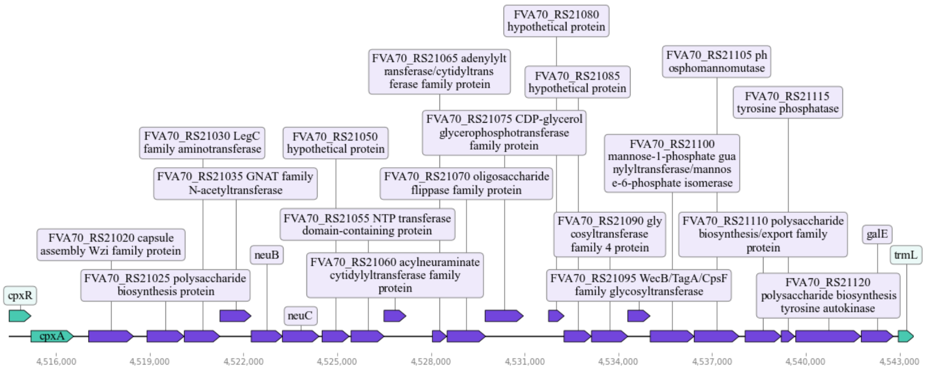
</td></tr>
<tr><td>Providencia sp. 1709051003</td><td>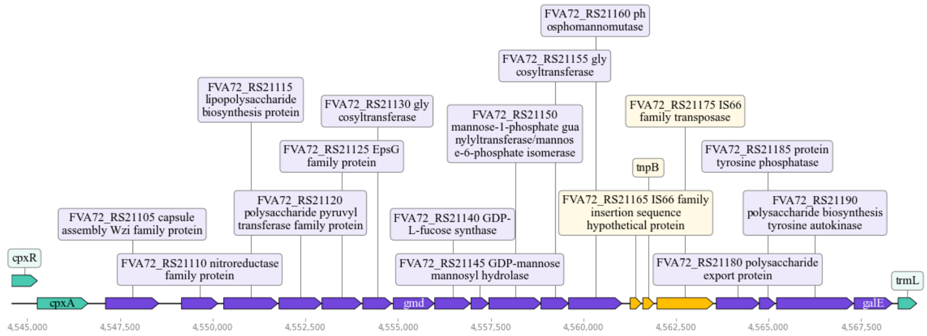
</td></tr>
<tr><td>Providencia rettgeri P01</td><td>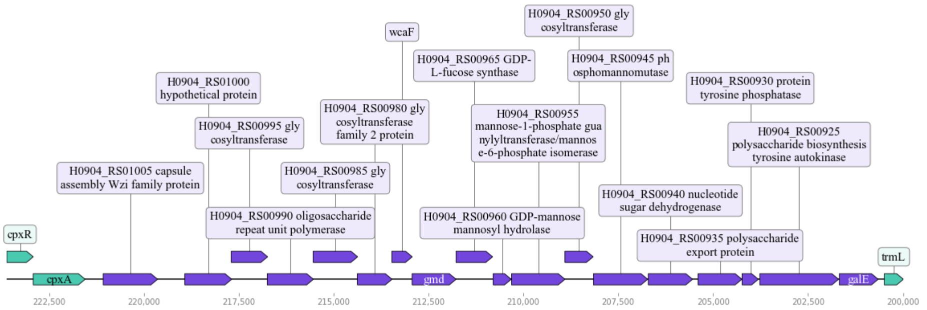
</td></tr>
<tr><td>Providencia rettgeri 2055</td><td>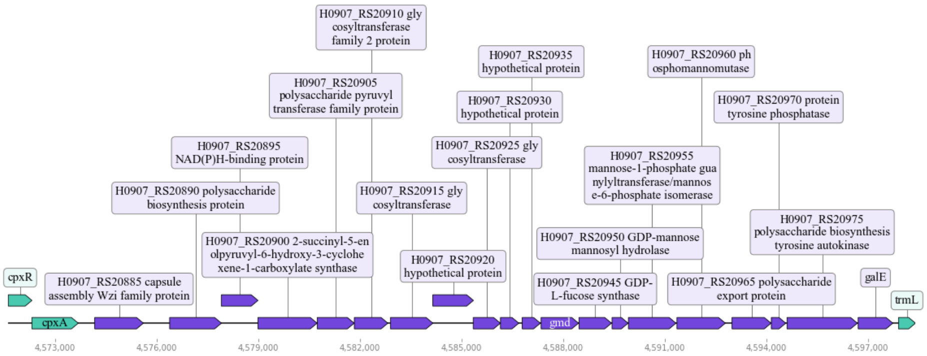
</td></tr>
</table>

**Table S4.2**. Composition of O-antigen operons based on PGAP annotation

<table>
<tr>
<td>Providencia alcalifaciens 1701003</td>
<td>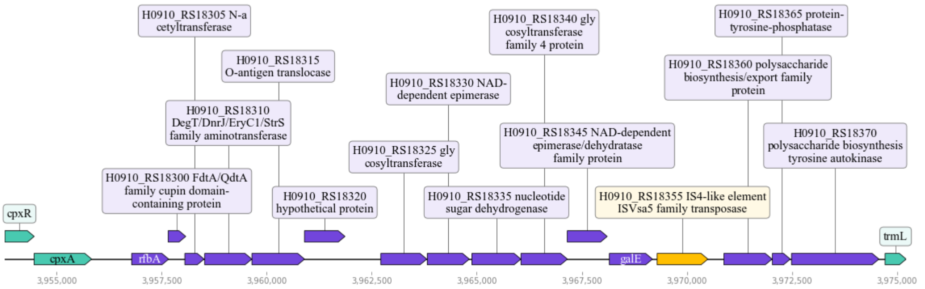
</td>
</tr>
<tr>
<td>Providencia rettgeri 2353217</td>
<td>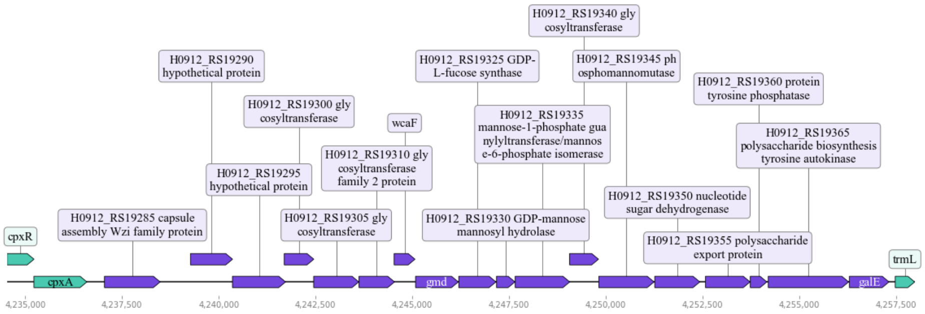
</td>
</tr>
<tr>
<td>Providencia rettgeri ZG2016M39-3</td>
<td>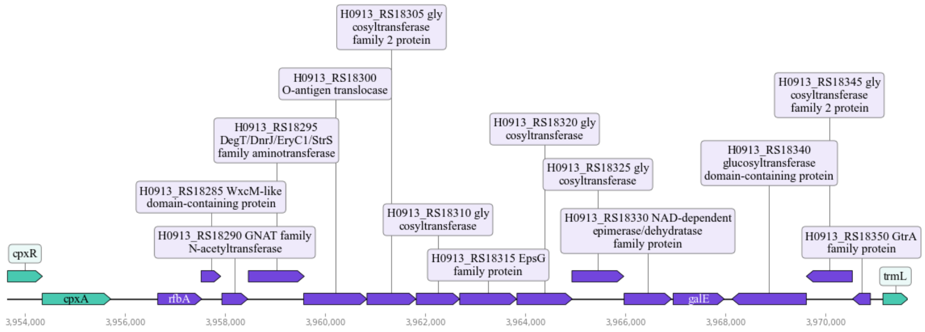
</td>
</tr>
<tr>
<td>Providencia sp. 2.29</td>
<td>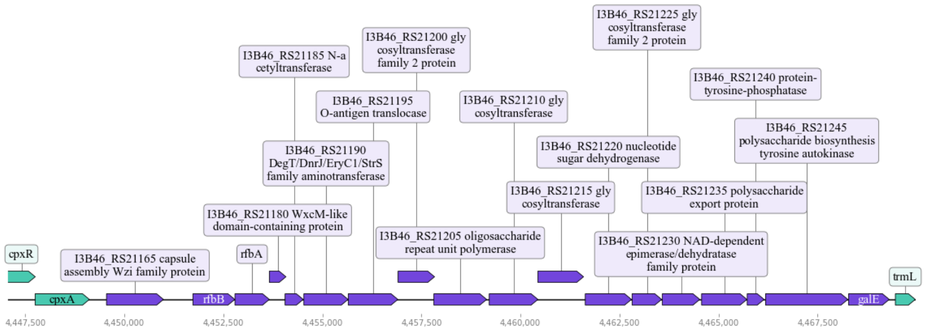
</td>
</tr>
</table>

**Table S4.2**. Composition of O-antigen operons based on PGAP annotation

<table>
<tr><td>

*Providencia rettgeri* W986

</td><td>
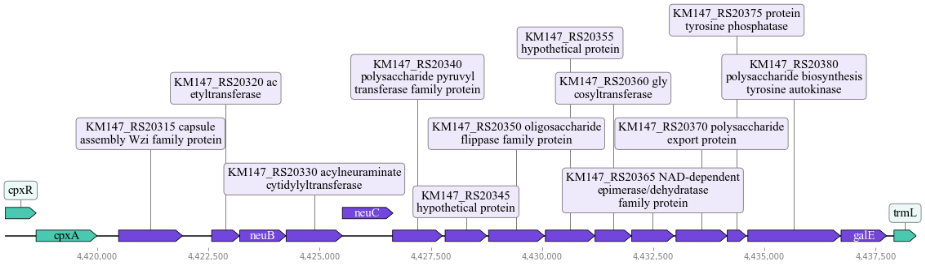

</td></tr>
<tr><td>

*Providencia rettgeri* PreM15973

</td><td>
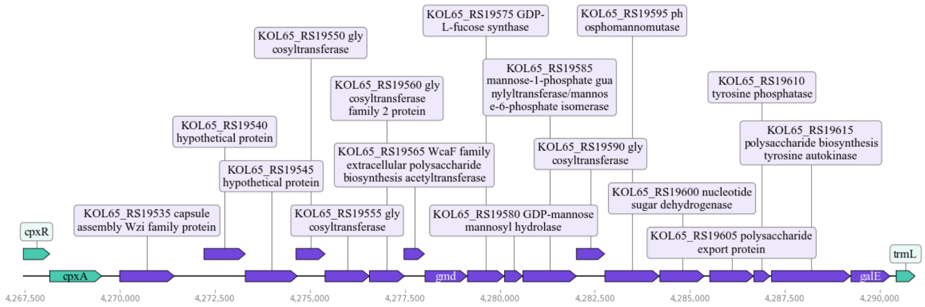

</td></tr>
<tr><td>

*Providencia rettgeri* PreM15628

</td><td>
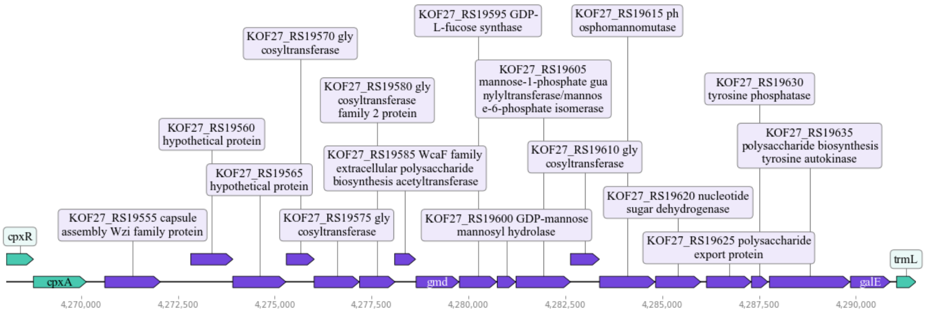

</td></tr>
<tr><td>

*Providencia rettgeri* PreM15758

</td><td>
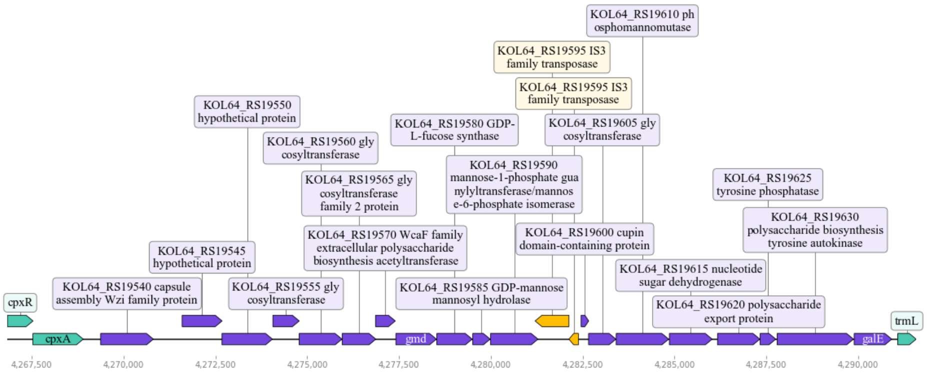

</td></tr>
<tr><td>

*Providencia rettgeri* FDAARGOS 1449

</td><td>
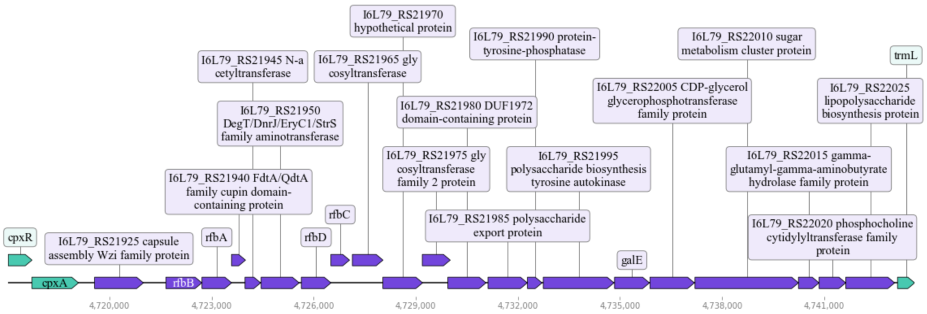

</td></tr>
</table>

**Table S4.2**. Composition of O-antigen operons based on PGAP annotation

<table>
<tr><td>Providencia rettgeri FDAARGOS 1451</td><td>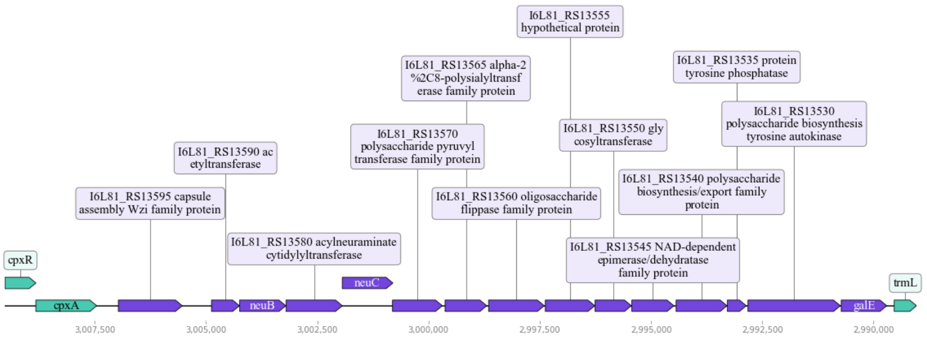
</td></tr>
<tr><td>Providencia heimbachae NCTC12003</td><td>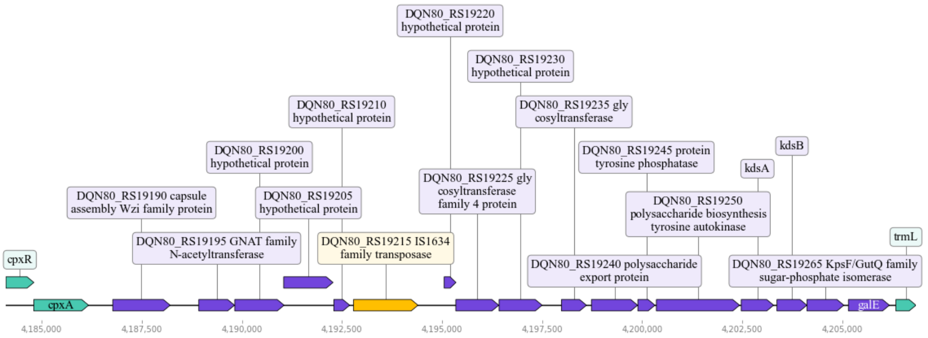
</td></tr>
<tr><td>Providencia rustigianii NCTC6933</td><td>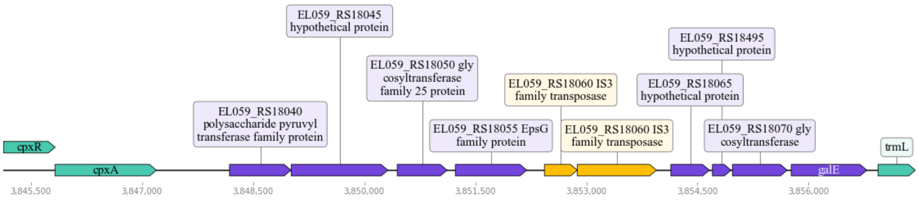
</td></tr>
<tr><td>Providencia rustigianii NCTC8113</td><td>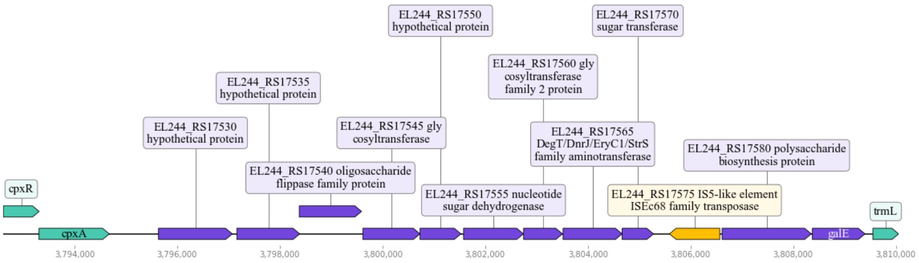
</td></tr>
</table>
