## Supplementary material for "Identification and comparison of somatic antigen composition for bacteria from *Providencia* genus": ./supplement/tableS4.pdf

**Table S5**. Gene phylogenetic trees used to clarify taxonomy classification of *Providencia* species used in the study

<table>
<tr><th>Gene name in Prokka annotation</th><th>Phylogenetic tree based on respective gene</th></tr>
<tr><td>wecA</td><td>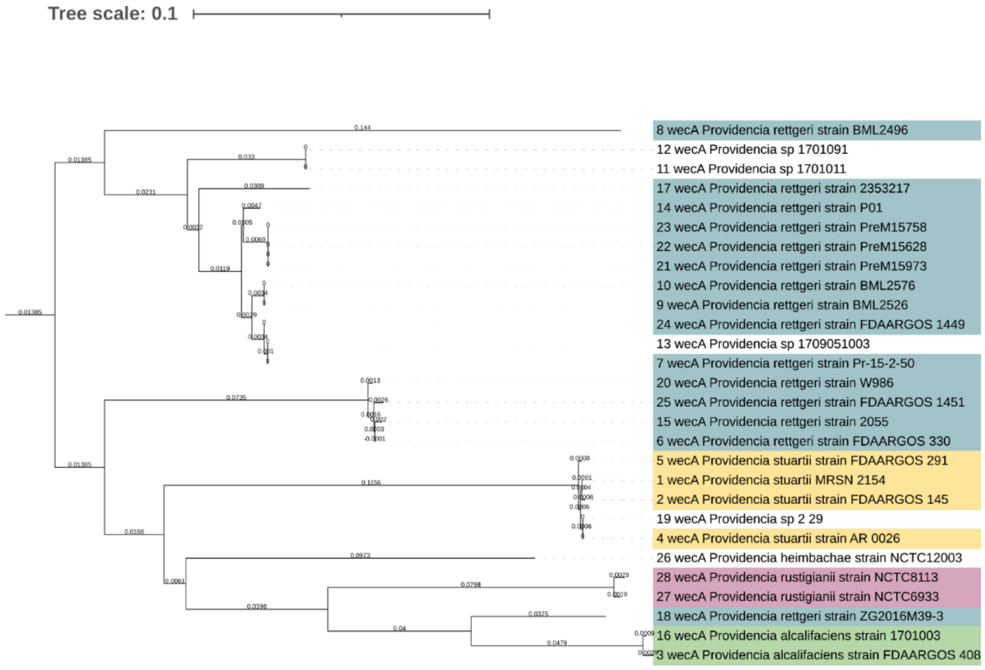
</td></tr>
<tr><td>wecB</td><td>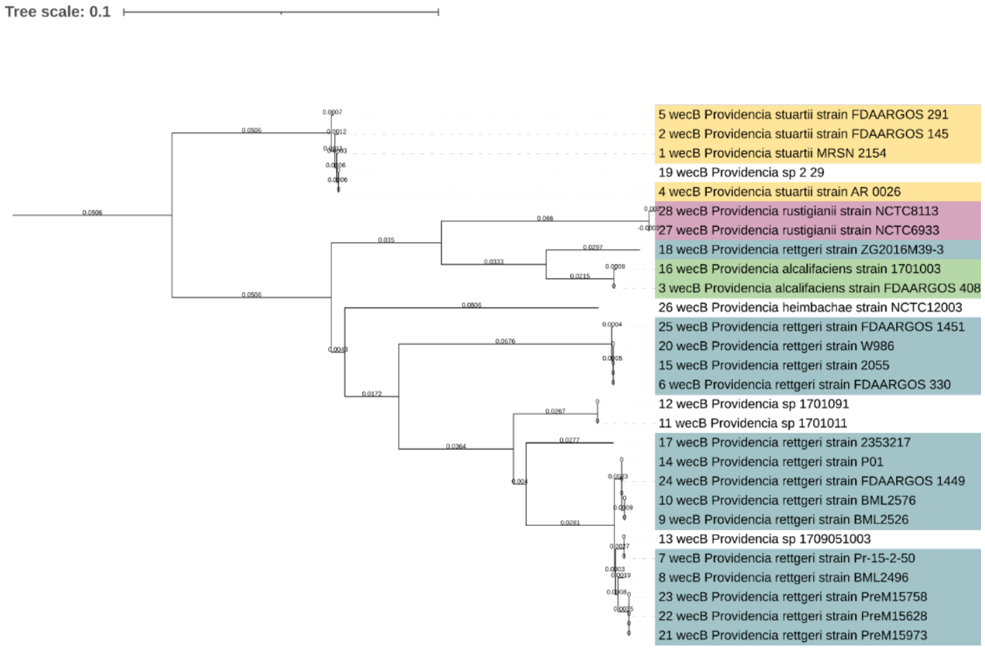
</td></tr>
</table>

**Table S5**. Gene phylogenetic trees used to clarify taxonomy classification of *Providencia* species used in the study

<table>
<tr>
<td>wecC</td>
<td>
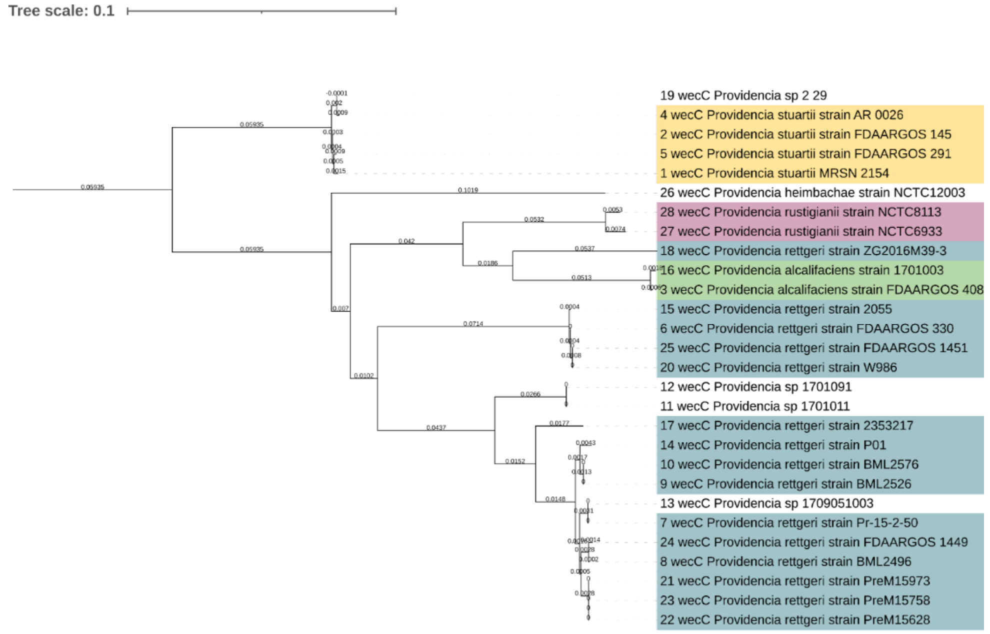

</td>
</tr>
<tr>
<td>wecD</td>
<td>
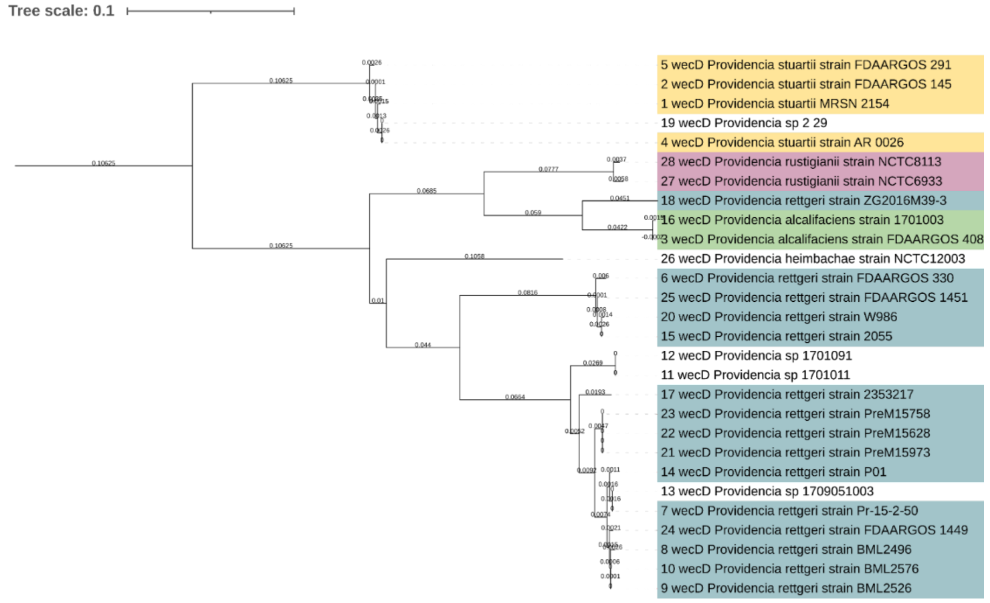

</td>
</tr>
</table>

**Table S5**. Gene phylogenetic trees used to clarify taxonomy classification of *Providencia* species used in the study

*wecE*

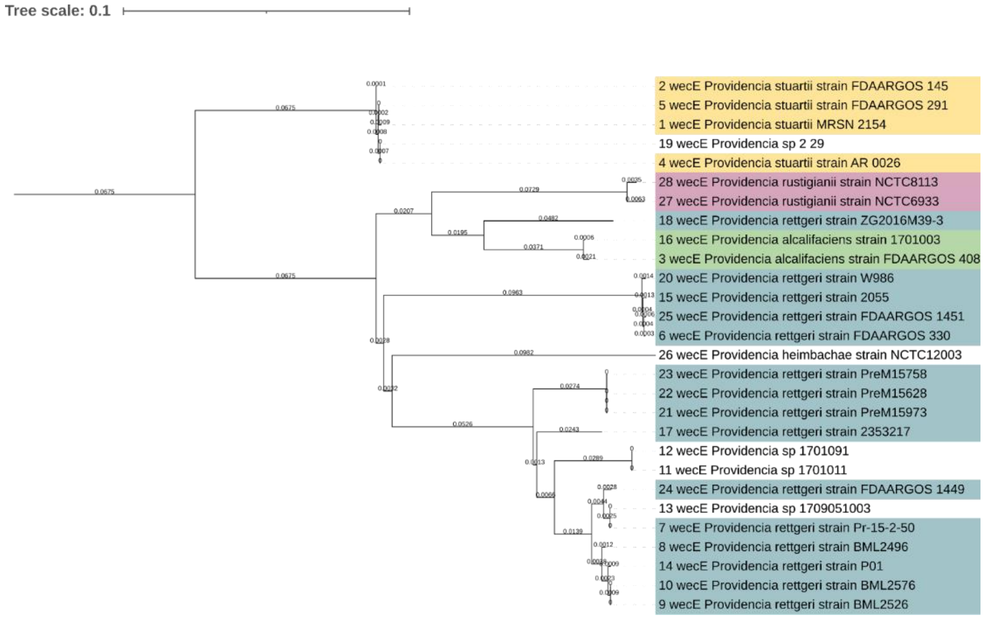


*wecF*

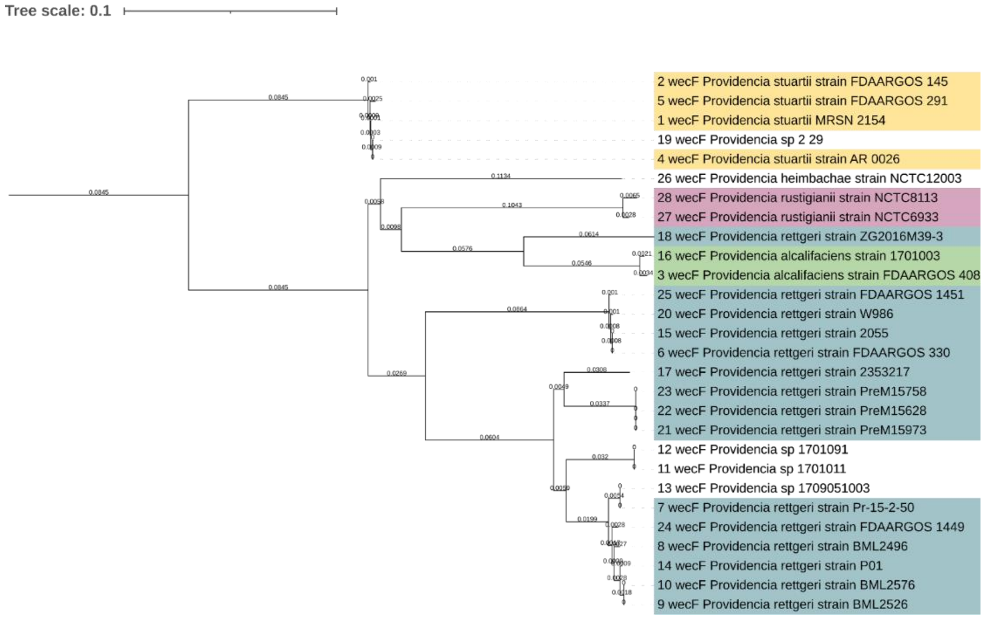

**Table S5**. Gene phylogenetic trees used to clarify taxonomy classification of *Providencia* species used in the study

<table>
<tr><td>wecG</td><td>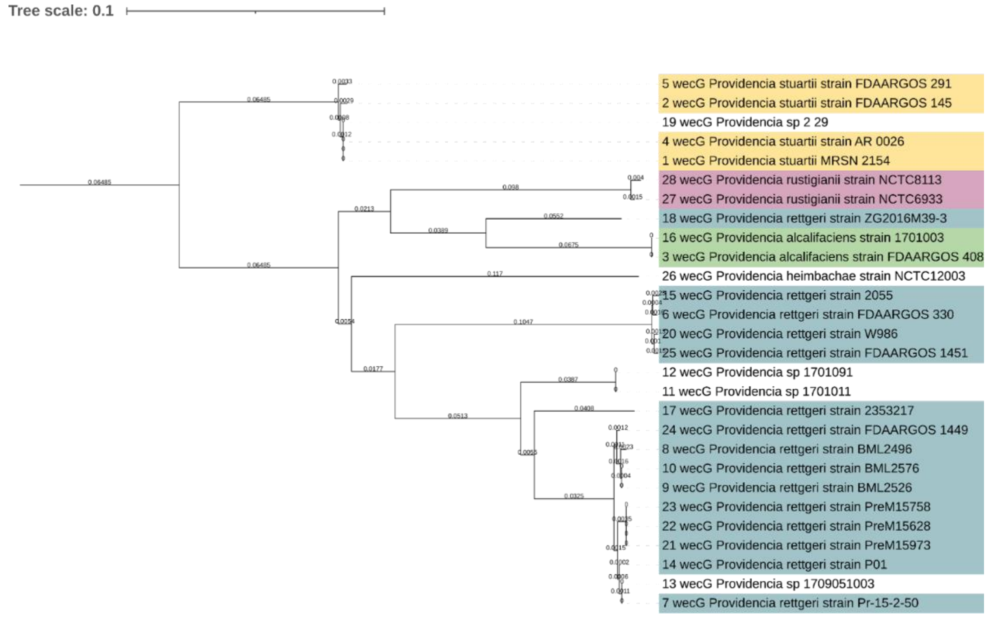
</td></tr>
<tr><td>rffH</td><td>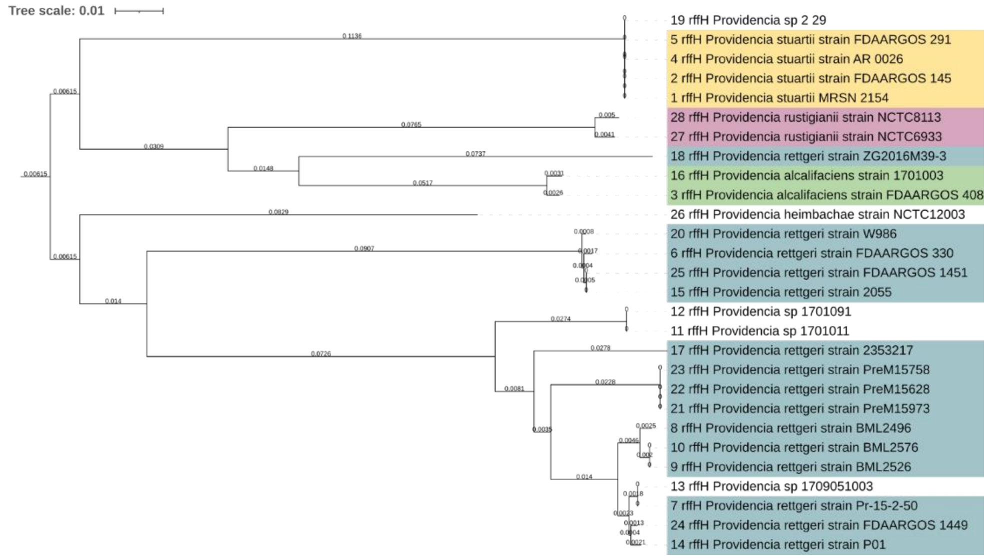
</td></tr>
<tr><td>rffG</td><td>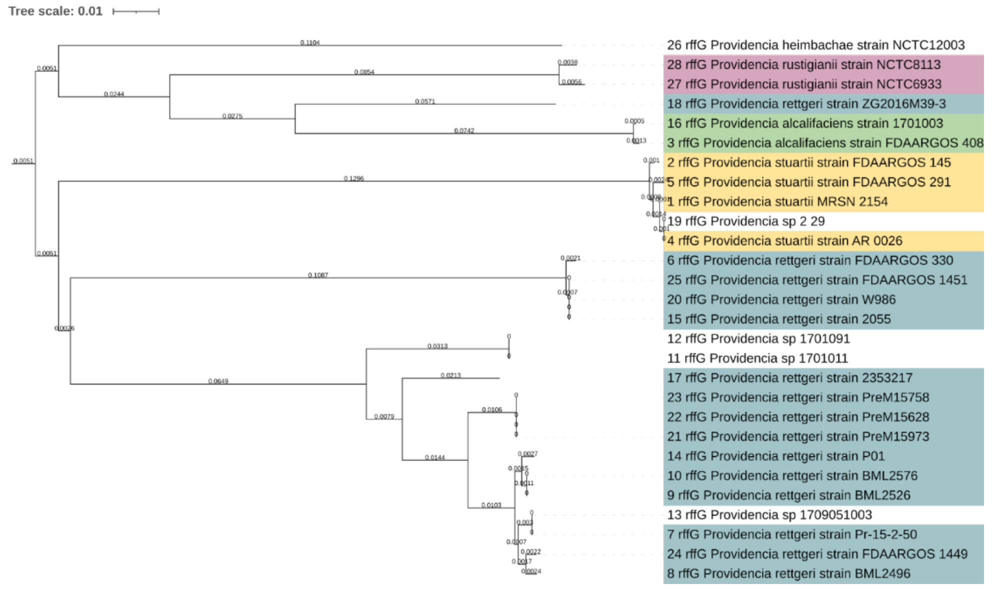
</td></tr>
</table>

**Table S5**. Gene phylogenetic trees used to clarify taxonomy classification of *Providencia* species used in the study

<table>
<tr>
<td>wzxE</td>
<td>
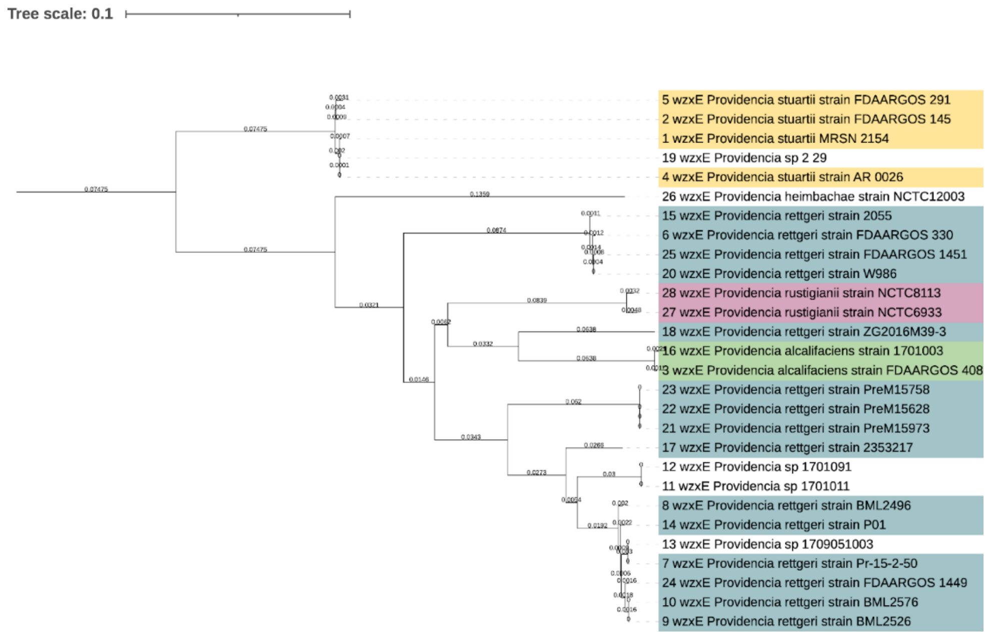

</td>
</tr>
<tr>
<td>wzzE</td>
<td>
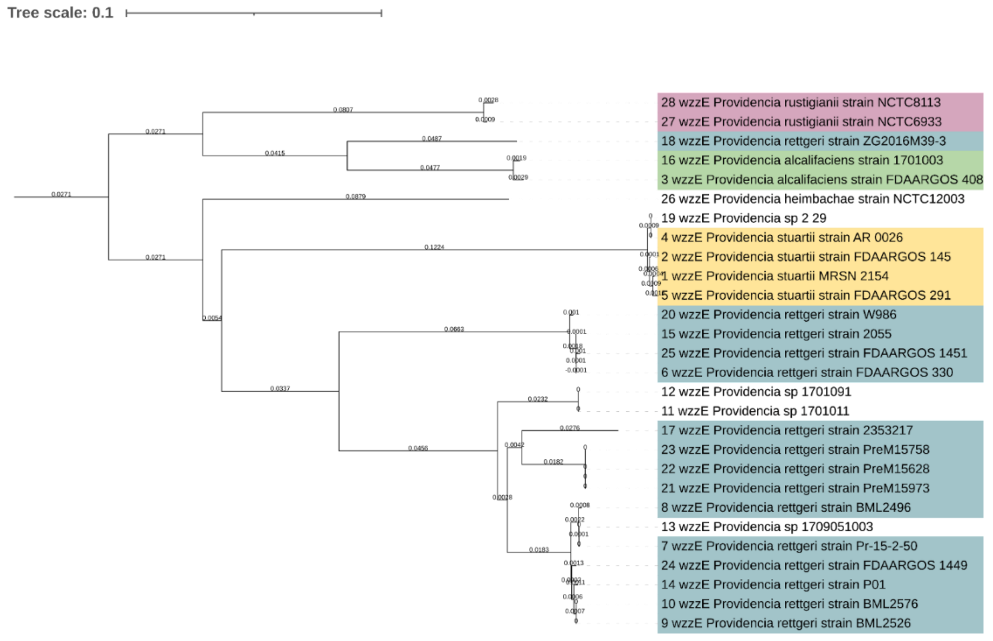

</td>
</tr>
</table>

**Table S5**. Gene phylogenetic trees used to clarify taxonomy classification of *Providencia* species used in the study

| | |
|---|---|
| *wzyE* | 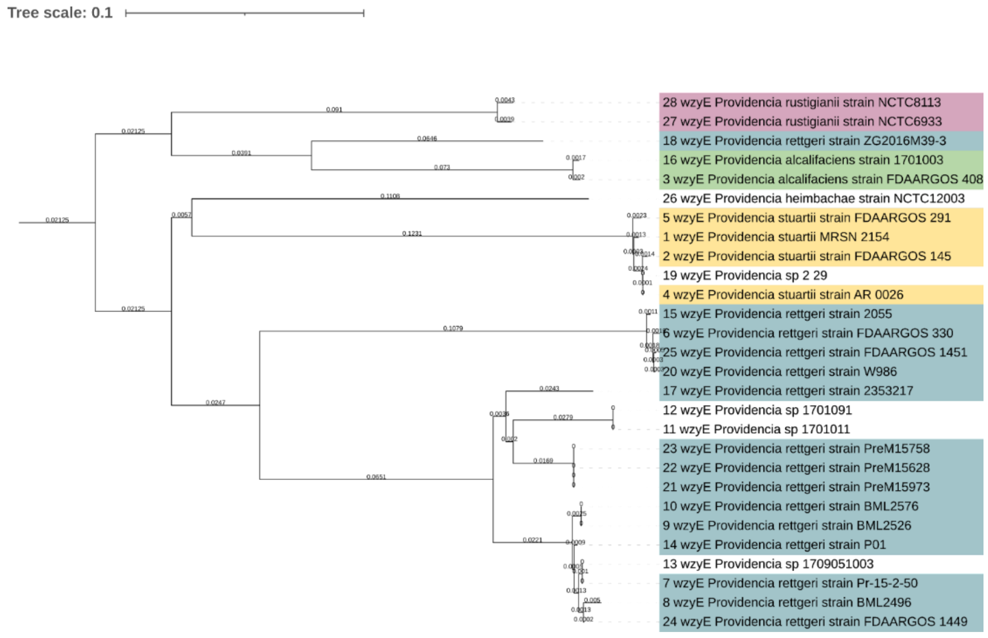 |
