## Supplementary material for "Identification and comparison of somatic antigen composition for bacteria from *Providencia* genus": ./supplement/tableS5.pdf

**Table S6**. Summary of genes identified within the O-antigen operon homologous to those from the Ovchinnikova et al. study. Gene homolog or its product name is provided according to Prokka annotation.

| **Genome ID** | ***Providencia* strain** | **Gene from Ovchinnikova et al.** | **Gene homolog name** | **Identity, %** |
|---|---|---|---|---|
| NZ_CP065420.1 | *Providencia sp.* 2.29 | *wzx* | *wzxE_2* | 71.31 |
| NZ_CP060726.1 | *P. rettgeri* strain YPR25 | *wzx* | *wzxE_2* | 75.54 |
| NZ_CP053896.1 | *Providencia rettgeri* strain YPR31 | *wzx* | *wzxE_2* | 75.54 |
| NZ_LS483422.1 | *P. heimbachae* strain NCTC12003 | *wzc* | *wzc_2* | 66.86 |
| NZ_CP077388.1 | *P. rettgeri* strain FDAARGOS 1451 | *wzc* | *wzc* | 66.89 |
| NZ_CP077317.1 | *P. rettgeri* strain FDAARGOS 1450 | *wzc* | *wzc_1* | 68.21 |
| NZ_CP077260.1 | *P. rettgeri* strain FDAARGOS 1449 | *wzc* | *wzc* | 68.88 |
| NZ_CP076407.1 | *P. rettgeri* strain PreM15758 | *wzc* | *wzc* | 65.18 |
| NZ_CP076406.1 | *P. rettgeri* strain PreM15973 | *wzc* | *wzc* | 65.18 |
| NZ_CP076405.1 | *P. rettgeri* strain PreM15628 | *wzc* | *wzc* | 65.18 |
| NZ_CP076258.1 | *P. rettgeri* strain W986 | *wzc* | *wzc* | 66.56 |
| NZ_CP066315.1 | *P. rettgeri* strain R39 | *wzc* | *wzc* | 66.56 |
| NZ_CP066071.1 | *P. stuartii* strain FDAARGOS_1040 | *wzc* | *wzc* | 66.62 |
| NZ_CP065420.1 | *Providencia* sp. 2.29 | *wzc* | *wzc* | 67.44 |
| NZ_CP062821.1 | *P. rettgeri* strain Res13-Sevr-LER2-35 | *wzc* | *wzc_2* | 69.02 |
| NZ_CP060726.1 | *P. rettgeri* strain YPR25 | *wzc* | *wzc* | 68.62 |
| NZ_CP059347.1 | *P. rettgeri* strain 2353217 | *wzc* | *wzc* | 69.17 |
| NZ_CP059346.1 | *P. alcalifaciens* strain 1701003 | *wzc* | *wzc* | 99.66 |

**Table S6**. Summary of genes identified within the O-antigen operon homologous to those from the Ovchinnikova et al. study. Gene homolog or its product name is provided according to Prokka annotation.

| | | | | |
|---|---|---|---|---|
| NZ_CP059345.1 | *P. rettgeri* strain 2055 | *wzc* | *wzc_2* | 67.14 |
| NZ_CP059298.1 | *P. rettgeri* strain P01 | *wzc* | *wzc_1* | 66.55 |
| NZ_CP058958.1 | *P. rettgeri* strain G0519 | *wzc* | *wzc* | 68.49 |
| NZ_CP053896.1 | *P. rettgeri* strain YPR31 | *wzc* | *wzc* | 68.62 |
| NZ_CP048796.1 | *P. vermicola* strain P8538 | *wzc* | *wzc* | 68.15 |
| NZ_CP048621.1 | *P. stuartii* strain MF1 | *wzc* | *wzc* | 63.57 |
| NZ_CP048621.1 | *P. stuartii* strain MF1 | *wzc* | *wzc* | 68.49 |
| NZ_CP044076.1 | *P. stuartii* strain FDAARGOS_645 | *wzc* | *wzc* | 66.47 |
| NZ_CP042861.1 | *Providencia* sp. 1709051003 | *wzc* | *wzc* | 65.28 |
| NZ_CP042860.1 | *Providencia* sp. 1701091 | *wzc* | *wzc* | 65.49 |
| NZ_CP042859.1 | *Providencia* sp. 1701011 | *wzc* | *wzc* | 65.49 |
| NZ_CP039844.1 | *P. rettgeri* strain Pr-15-2-50 | *wzc* | *wzc* | 65.28 |
| NZ_CP031123.2 | *P. huaxiensis* strain WCHPr000369 | *wzc* | *wzc* | 61.61 |
| NZ_CP031123.2 | *P. huaxiensis* strain WCHPr000369 | *wzc* | *wzc* | 67.56 |
| NZ_CP029736.1 | *P. rettgeri* strain AR_0082 | *wzc* | *wzc* | 66.26 |
| NZ_CP028384.1 | *P. heimbachae* strain 99101 | *wzc* | *wzc* | 66.5 |
| NZ_CP027418.1 | *P. rettgeri* strain FDAARGOS_330 | *wzc* | *wzc* | 66.91 |
| NZ_CP027398.1 | *P. stuartii* strain FDAARGOS_291 | *wzc* | *wzc* | 70.88 |
| NZ_CP027398.1 | *P. stuartii* strain FDAARGOS_291 | *wzc* | *wzc* | 68.86 |
| NZ_CP026704.1 | *P. stuartii* strain AR_0026 | *wzc* | *wzc* | 66.89 |

**Table S6**. Summary of genes identified within the O-antigen operon homologous to those from the Ovchinnikova et al. study. Gene homolog or its product name is provided according to Prokka annotation.

| | | | | |
|---|---|---|---|---|
| NZ_CP023536.1 | *P. alcalifaciens* strain FDAARGOS_408 | *wzc* | *wzc* | 98.56 |
| NZ_CP017671.1 | *P. rettgeri* strain RB151 | *wzc* | *wzc_2* | 66.91 |
| NZ_CP017054.1 | *P. stuartii* strain BE2467 | *wzc* | *wzc* | 66.57 |
| NZ_CP014024.2 | *P. stuartii* strain FDAARGOS_145 | *wzc* | *wzc* | 70.81 |
| NZ_CP008920.1 | *P. stuartii* strain ATCC 33672 | *wzc* | *wzc* | 66.62 |
| NZ_AP022375.1 | *P. rettgeri* strain BML2576 | *wzc* | *wzc* | 68.17 |
| NZ_AP022374.1 | *P. stuartii* strain BML2537 | *wzc* | *wzc* | 67.48 |
| NZ_AP022373.1 | *P. rettgeri* strain BML2531 | *wzc* | *wzc* | 68.56 |
| NZ_AP022372.1 | *P. rettgeri* strain BML2526 | *wzc* | *wzc* | 68.17 |
| NZ_AP022371.1 | *P. rettgeri* strain BML2496 | *wzc* | *wzc* | 70.76 |
| NZ_LS483422.1 | *P. heimbachae* strain NCTC12003 | *wzb* | *wzb_2* | 72.49 |
| NZ_CP077388.1 | *P. rettgeri* strain FDAARGOS 1451 | *wzb* | *wzb* | 69.21 |
| NZ_CP077317.1 | *P. rettgeri* strain FDAARGOS 1450 | *wzb* | *wzb* | 69.66 |
| NZ_CP077260.1 | *P. rettgeri* strain FDAARGOS 1449 | *wzb* | *wzb* | 71.3 |
| NZ_CP076407.1 | *P. rettgeri* strain PreM15758 | *wzb* | *wzb* | 67.82 |
| NZ_CP076406.1 | *P. rettgeri* strain PreM15973 | *wzb* | *wzb* | 67.82 |
| NZ_CP076405.1 | *P. rettgeri* strain PreM15628 | *wzb* | *wzb* | 67.82 |
| NZ_CP076258.1 | *P. rettgeri* strain W986 | *wzb* | *wzb* | 69.21 |
| NZ_CP066315.1 | *P. rettgeri* strain R39 | *wzb* | *wzb* | 69.21 |

**Table S6**. Summary of genes identified within the O-antigen operon homologous to those from the Ovchinnikova et al. study. Gene homolog or its product name is provided according to Prokka annotation.

| | | | | |
|---|---|---|---|---|
| NZ_CP066071.1 | *P. stuartii* strain FDAARGOS_1040 | *wzb* | *wzb* | 73.89 |
| NZ_CP065420.1 | *Providencia* sp. 2.29 | *wzb* | *wzb* | 73.43 |
| NZ_CP062821.1 | *P. rettgeri* strain Res13-Sevr-LER2-35 | *wzb* | *wzb_2* | 68.82 |
| NZ_CP060726.1 | *P. rettgeri* strain YPR25 | *wzb* | *wzb* | 68.51 |
| NZ_CP059347.1 | *P. rettgeri* strain 2353217 | *wzb* | *wzb* | 69.05 |
| NZ_CP059346.1 | *P. alcalifaciens* strain 1701003 | *wzb* | *wzb* | 99.77 |
| NZ_CP059345.1 | *P. rettgeri* strain 2055 | *wzb* | *wzb_2* | 68.51 |
| NZ_CP059298.1 | *P. rettgeri* strain P01 | *wzb* | *wzb_1* | 68.98 |
| NZ_CP058958.1 | *P. rettgeri* strain G0519 | *wzb* | *wzb* | 68.51 |
| NZ_CP053896.1 | *P. rettgeri* strain YPR31 | *wzb* | *wzb* | 68.51 |
| NZ_CP048796.1 | *P. vermicola* strain P8538 | *wzb* | *wzb* | 74.36 |
| NZ_CP048621.1 | *P. stuartii* strain MF1 | *wzb* | *wzb* | 78.37 |
| NZ_CP044076.1 | *P. stuartii* strain FDAARGOS_645 | *wzb* | *wzb* | 72.29 |
| NZ_CP042861.1 | *Providencia* sp. 1709051003 | *wzb* | *wzb* | 68.28 |
| NZ_CP042860.1 | *Providencia* sp. 1701091 | *wzb* | *wzb* | 67.36 |
| NZ_CP042859.1 | *Providencia* sp. 1701011 | *wzb* | *wzb* | 67.36 |
| NZ_CP039844.1 | *P. rettgeri* strain Pr-15-2-50 | *wzb* | *wzb* | 68.28 |
| NZ_CP031123.2 | *P. huaxiensis* strain WCHPr000369 | *wzb* | *wzb* | 71.67 |
| NZ_CP029736.1 | *P. rettgeri* strain AR_0082 | *wzb* | *wzb* | 68.29 |
| NZ_CP028384.1 | *P. heimbachae* strain 99101 | *wzb* | *wzb* | 71.1 |

**Table S6**. Summary of genes identified within the O-antigen operon homologous to those from the Ovchinnikova et al. study. Gene homolog or its product name is provided according to Prokka annotation.

| | | | | |
|---|---|---|---|---|
| NZ_CP027418.1 | *P. rettgeri* strain FDAARGOS_330 | *wzb* | *wzb* | 70.9 |
| NZ_CP027398.1 | *P. stuartii* strain FDAARGOS_291 | *wzb* | *wzb* | 73.89 |
| NZ_CP026704.1 | *P. stuartii* strain AR_0026 | *wzb* | *wzb* | 70.44 |
| NZ_CP023536.1 | *P. alcalifaciens* strain FDAARGOS_408 | *wzb* | *wzb* | 97.9 |
| NZ_CP017671.1 | *P. rettgeri* strain RB151 | *wzb* | *wzb_2* | 70.9 |
| NZ_CP017054.1 | *P. stuartii* strain BE2467 | *wzb* | *wzb* | 72.29 |
| NZ_CP014024.2 | *P. stuartii* strain FDAARGOS_145 | *wzb* | *wzb* | 72.22 |
| NZ_CP008920.1 | *P. stuartii* strain ATCC 33672 | *wzb* | *wzb* | 73.89 |
| NZ_AP022375.1 | *P. rettgeri* strain BML2576 | *wzb* | *wzb* | 68.13 |
| NZ_AP022374.1 | *P. stuartii* strain BML2537 | *wzb* | *wzb* | 71.99 |
| NZ_AP022373.1 | *P. rettgeri* strain BML2531 | *wzb* | *wzb* | 68.13 |
| NZ_AP022372.1 | *P. rettgeri* strain BML2526 | *wzb* | *wzb* | 68.13 |
| NZ_AP022371.1 | *P. rettgeri* strain BML2496 | *wzb* | *wzb* | 66.9 |
| NZ_LS483422.1 | *P. heimbachae* strain NCTC12003 | *wza* | hypothetical protein | 72.45 |
| NZ_CP077388.1 | *P. rettgeri* strain FDAARGOS 1451 | *wza* | hypothetical protein | 70.99 |
| NZ_CP077317.1 | *P. rettgeri* strain FDAARGOS 1450 | *wza* | hypothetical protein | 74.15 |
| NZ_CP077260.1 | *P. rettgeri* strain FDAARGOS 1449 | *wza* | hypothetical protein | 71.95 |
| NZ_CP076407.1 | *P. rettgeri* strain PreM15758 | *wza* | hypothetical protein | 72.18 |
| NZ_CP076406.1 | *P. rettgeri* strain PreM15973 | *wza* | hypothetical protein | 72.18 |

**Table S6**. Summary of genes identified within the O-antigen operon homologous to those from the Ovchinnikova et al. study. Gene homolog or its product name is provided according to Prokka annotation.

| | | | | |
|---|---|---|---|---|
| NZ_CP076405.1 | *P. rettgeri* strain PreM15628 | *wza* | hypothetical protein | 72.18 |
| NZ_CP076258.1 | *P. rettgeri* strain W986 | *wza* | hypothetical protein | 71.62 |
| NZ_CP066315.1 | *P. rettgeri* strain R39 | *wza* | hypothetical protein | 71.62 |
| NZ_CP066071.1 | *P. stuartii* strain FDAARGOS_1040 | *wza* | hypothetical protein | 74.03 |
| NZ_CP065420.1 | *Providencia* sp. 2.29 | *wza* | hypothetical protein | 73.63 |
| NZ_CP062821.1 | *P. rettgeri* strain Res13-Sevr-LER2-35 | *wza* | hypothetical protein | 72.5 |
| NZ_CP060726.1 | *P. rettgeri* strain YPR25 | *wza* | hypothetical protein | 71.32 |
| NZ_CP059347.1 | *P. rettgeri* strain 2353217 | *wza* | hypothetical protein | 72.55 |
| NZ_CP059346.1 | *P. alcalifaciens* strain 1701003 | *wza* | hypothetical protein | 99.04 |
| NZ_CP059345.1 | *P. rettgeri* strain 2055 | *wza* | hypothetical protein | 70.27 |
| NZ_CP059298.1 | *P. rettgeri* strain P01 | *wza* | hypothetical protein | 72.27 |
| NZ_CP058958.1 | *P. rettgeri* strain G0519 | *wza* | hypothetical protein | 71.72 |
| NZ_CP053896.1 | *P. rettgeri* strain YPR31 | *wza* | hypothetical protein | 71.32 |
| NZ_CP048796.1 | *P. vermicola* strain P8538 | *wza* | hypothetical protein | 72.87 |
| NZ_CP048621.1 | *P. stuartii* strain MF1 | *wza* | hypothetical protein | 74.19 |
| NZ_CP044076.1 | *P. stuartii* strain FDAARGOS_645 | *wza* | hypothetical protein | 73.82 |
| NZ_CP042861.1 | *Providencia* sp. 1709051003 | *wza* | hypothetical protein | 70.96 |
| NZ_CP042860.1 | *Providencia* sp. 1701091 | *wza* | hypothetical protein | 72.34 |
| NZ_CP042859.1 | *Providencia* sp. 1701011 | *wza* | hypothetical protein | 72.34 |

**Table S6**.  Summary of genes identified within the O-antigen operon homologous to those from the Ovchinnikova et al. study. Gene homolog or its product name is provided according to Prokka annotation.

| | | | | |
|---|---|---|---|---|
| NZ_CP039844.1 | *P. rettgeri* strain Pr-15-2-50 | *wza* | hypothetical protein | 70.96 |
| NZ_CP031123.2 | *P. huaxiensis* strain WCHPr000369 | *wza* | hypothetical protein | 71.52 |
| NZ_CP029736.1 | *P. rettgeri* strain AR_0082 | *wza* | hypothetical protein | 72.39 |
| NZ_CP028384.1 | *P. heimbachae* strain 99101 | *wza* | hypothetical protein | 72.3 |
| NZ_CP027418.1 | *P. rettgeri* strain FDAARGOS_330 | *wza* | hypothetical protein | 71.49 |
| NZ_CP027398.1 | *P. stuartii* strain FDAARGOS_291 | *wza* | hypothetical protein | 73.79 |
| NZ_CP026704.1 | *P. stuartii* strain AR_0026 | *wza* | hypothetical protein | 73.11 |
| NZ_CP023536.1 | *P. alcalifaciens* strain FDAARGOS_408 | *wza* | hypothetical protein | 98.52 |
| NZ_CP017671.1 | *P. rettgeri* strain RB151 | *wza* | hypothetical protein | 71.49 |
| NZ_CP017054.1 | *P. stuartii* strain BE2467 | *wza* | hypothetical protein | 73.92 |
| NZ_CP014024.2 | *P. stuartii* strain FDAARGOS_145 | *wza* | hypothetical protein | 74.1 |
| NZ_CP008920.1 | *P. stuartii* strain ATCC 33672 | *wza* | hypothetical protein | 74.03 |
| NZ_AP022375.1 | *P. rettgeri* strain BML2576 | *wza* | hypothetical protein | 71.83 |
| NZ_AP022374.1 | *P. stuartii* strain BML2537 | *wza* | hypothetical protein | 73.01 |
| NZ_AP022373.1 | *P. rettgeri* strain BML2531 | *wza* | hypothetical protein | 72.53 |
| NZ_AP022372.1 | *P. rettgeri* strain BML2526 | *wza* | hypothetical protein | 71.83 |
| NZ_AP022371.1 | *P. rettgeri* strain BML2496 | *wza* | hypothetical protein | 72.01 |
| NZ_CP065420.1 | *Providencia* sp. 2.29 | *wpaD* | *tuaG* | 72.78 |
| NZ_CP060726.1 | *P. rettgeri* strain YPR25 | *wpaD* | *tuaG* | 66.03 |

**Table S6**. Summary of genes identified within the O-antigen operon homologous to those from the Ovchinnikova et al. study. Gene homolog or its product name is provided according to Prokka annotation.

| | | | | |
|---|---|---|---|---|
| NZ_CP059298.1 | *P. rettgeri* strain P01 | *wpaD* | *ugd_1* | 86.67 |
| NZ_CP053896.1 | *P. rettgeri* strain YPR31 | *wpaD* | *tuaG* | 66.03 |
| NZ_AP022375.1 | *P. rettgeri* strain BML2576 | *wpaD* | *tuaG* | 73.73 |
| NZ_AP022372.1 | *P. rettgeri* strain BML2526 | *wpaD* | *tuaG* | 73.73 |
| NZ_AP022375.1 | *P. rettgeri* strain BML2576 | *wpaB* | *mshA* | 67.85 |
| NZ_AP022372.1 | *P. rettgeri* strain BML2526 | *wpaB* | *mshA* | 67.85 |
| NZ_CP060726.1 | *P. rettgeri* strain YPR25 | *wpaA* | hypothetical protein | 65.41 |
| NZ_CP053896.1 | *P. rettgeri* strain YPR31 | *wpaA* | hypothetical protein | 65.41 |
| NZ_LR134396.1 | *P. rustigianii* strain NCTC8113 | *ugd* | *ugd_2* | 85.95 |
| NZ_CP076407.1 | *P. rettgeri* strain PreM15758 | *ugd* | *ugd_1* | 83.03 |
| NZ_CP076406.1 | *P. rettgeri* strain PreM15973 | *ugd* | *ugd_1* | 83.03 |
| NZ_CP076405.1 | *P. rettgeri* strain PreM15628 | *ugd* | *ugd_1* | 83.03 |
| NZ_CP066071.1 | *P. stuartii* strain FDAARGOS_1040 | *ugd* | *ugd_1* | 81.58 |
| NZ_CP065420.1 | *Providencia* sp. 2.29 | *ugd* | *ugd_1* | 81.75 |
| NZ_CP060726.1 | *P. rettgeri* strain YPR25 | *ugd* | *ugd_1* | 82.16 |
| NZ_CP059347.1 | *P. rettgeri* strain 2353217 | *ugd* | *ugd_1* | 81.23 |
| NZ_CP059346.1 | *P. alcalifaciens* strain 1701003 | *ugd* | *ugd_1* | 98.11 |
| NZ_CP059298.1 | *P. rettgeri* strain P01 | *ugd* | *ugd_1* | 81.58 |
| NZ_CP053896.1 | *P. rettgeri* strain YPR31 | *ugd* | *ugd_1* | 82.16 |
| NZ_CP044076.1 | *P. stuartii* strain FDAARGOS_645 | *ugd* | *ugd_2* | 81.65 |

**Table S6**. Summary of genes identified within the O-antigen operon homologous to those from the Ovchinnikova et al. study. Gene homolog or its product name is provided according to Prokka annotation.

| | | | | |
|---|---|---|---|---|
| NZ_CP028384.1 | *P. heimbachae* strain 99101 | *ugd* | *ugd_1* | 81.49 |
| NZ_CP027398.1 | *P. stuartii* strain FDAARGOS_291 | *ugd* | *ugd_1* | 81.65 |
| NZ_CP017054.1 | *P. stuartii* strain BE2467 | *ugd* | *ugd_2* | 81.65 |
| NZ_CP014024.2 | *P. stuartii* strain FDAARGOS_145 | *ugd* | *ugd_1* | 81.83 |
| NZ_CP008920.1 | *P. stuartii* strain ATCC 33672 | *ugd* | *ugd_1* | 81.58 |
| NZ_AP022375.1 | *P. rettgeri* strain BML2576 | *ugd* | *ugd_1* | 82.25 |
| NZ_AP022373.1 | *P. rettgeri* strain BML2531 | *ugd* | *ugd_1* | 81.15 |
| NZ_AP022372.1 | *P. rettgeri* strain BML2526 | *ugd* | *ugd_1* | 82.25 |
| NZ_AP022371.1 | *P. rettgeri* strain BML2496 | *ugd* | *ugd_1* | 82.26 |
| NZ_CP077317.1 | *P. rettgeri* strain FDAARGOS 1450 | *rmlA* | *rmlA* | 66.19 |
| NZ_CP077260.1 | *P. rettgeri* strain FDAARGOS 1449 | *rmlA* | *rffH_2* | 80.78 |
| NZ_CP066071.1 | *P. stuartii* strain FDAARGOS_1040 | *rmlA* | *rffH_2* | 73.65 |
| NZ_CP065420.1 | *Providencia* sp. 2.29 | *rmlA* | *rffH_2* | 80.63 |
| NZ_CP060726.1 | *P. rettgeri* strain YPR25 | *rmlA* | *rffH_2* | 82.74 |
| NZ_CP059348.1 | *P. rettgeri* strain ZG2016M39-3 | *rmlA* | *rffH_2* | 90.59 |
| NZ_CP059346.1 | *P. alcalifaciens* strain 1701003 | *rmlA* | *rffH_2* | 97.05 |
| NZ_CP053896.1 | *P. rettgeri* strain YPR31 | *rmlA* | *rffH_2* | 82.74 |
| NZ_CP029736.1 | *P. rettgeri* strain AR_0082 | *rmlA* | *rffH_1* | 81.43 |
| NZ_CP023536.1 | *P. alcalifaciens* strain FDAARGOS_408 | *rmlA* | *rffH_1* | 97.05 |

**Table S6**. Summary of genes identified within the O-antigen operon homologous to those from the Ovchinnikova et al. study. Gene homolog or its product name is provided according to Prokka annotation.

| | | | | |
|---|---|---|---|---|
| NZ_CP014024.2 | *P. stuartii* strain FDAARGOS_145 | *rmlA* | *rffH_2* | 73.41 |
| NZ_CP008920.1 | *P. stuartii* strain ATCC 33672 | *rmlA* | *rffH_2* | 73.65 |
| NZ_AP022374.1 | *P. stuartii* strain BML2537 | *rmlA* | *rffH_2* | 73.61 |
| NZ_AP022373.1 | *P. rettgeri* strain BML2531 | *rmlA* | *rmlA* | 66.19 |
| NZ_CP077260.1 | *P. rettgeri* strain FDAARGOS 1449 | *qdtB* | *fdtB* | 72.98 |
| NZ_CP065420.1 | *Providencia* sp. 2.29 | *qdtB* | *fdtB* | 73.85 |
| NZ_CP060726.1 | *P. rettgeri* strain YPR25 | *qdtB* | *fdtB* | 76.33 |
| NZ_CP059348.1 | *P. rettgeri* strain ZG2016M39-3 | *qdtB* | *fdtB* | 85.24 |
| NZ_CP059346.1 | *P. alcalifaciens* strain 1701003 | *qdtB* | *fdtB* | 85.16 |
| NZ_CP053896.1 | *P. rettgeri* strain YPR31 | *qdtB* | *fdtB* | 76.33 |
| NZ_CP029736.1 | *P. rettgeri* strain AR_0082 | *qdtB* | *fdtB* | 73.18 |
| NZ_CP023536.1 | *P. alcalifaciens* strain FDAARGOS_408 | *qdtB* | *fdtB* | 84.59 |
| NZ_CP077260.1 | *P. rettgeri* strain FDAARGOS 1449 | *qdtA* | *fdtA* | 77.12 |
| NZ_CP065420.1 | *Providencia* sp. 2.29 | *qdtA* | *fdtA* | 80.58 |
| NZ_CP060726.1 | *P. rettgeri* strain YPR25 | *qdtA* | *fdtA* | 79.22 |
| NZ_CP059348.1 | *P. rettgeri* strain ZG2016M39-3 | *qdtA* | *fdtA* | 78.2 |
| NZ_CP059346.1 | *P. alcalifaciens* strain 1701003 | *qdtA* | *fdtA* | 79.18 |
| NZ_CP053896.1 | *P. rettgeri* strain YPR31 | *qdtA* | *fdtA* | 79.22 |
| NZ_CP029736.1 | *P. rettgeri* strain AR_0082 | *qdtA* | *fdtA* | 81.5 |

**Table S6**. Summary of genes identified within the O-antigen operon homologous to those from the Ovchinnikova et al. study. Gene homolog or its product name is provided according to Prokka annotation.

| | | | | |
|---|---|---|---|---|
| NZ_CP023536.1 | *P. alcalifaciens* strain FDAARGOS_408 | *qdtA* | *fdtA* | 79.3 |
| NZ_CP077388.1 | *P. rettgeri* strain FDAARGOS 1451 | *gne* | *gnu* | 83.33 |
| NZ_CP076258.1 | *P. rettgeri* strain W986 | *gne* | *gnu* | 82.91 |
| NZ_CP066315.1 | *P. rettgeri* strain R39 | *gne* | *gnu* | 82.91 |
| NZ_CP065420.1 | *Providencia* sp. 2.29 | *gne* | *gnu* | 80.06 |
| NZ_CP062821.1 | *P. rettgeri* strain Res13-Sevr-LER2-35 | *gne* | *gnu* | 82.49 |
| NZ_CP059348.1 | *P. rettgeri* strain ZG2016M39-3 | *gne* | *gnu* | 93.67 |
| NZ_CP059346.1 | *P. alcalifaciens* strain 1701003 | *gne* | *gnu* | 97.78 |
| NZ_CP031123.2 | *P. huaxiensis* strain WCHPr000369 | *gne* | *gnu* | 81.47 |
| NZ_CP029736.1 | *P. rettgeri* strain AR_0082 | *gne* | *gnu* | 82.91 |
| NZ_CP027418.1 | *P. rettgeri* strain FDAARGOS_330 | *gne* | *gnu* | 83.23 |
| NZ_CP026704.1 | *P. stuartii* strain AR_0026 | *gne* | *gnu* | 81.24 |
| NZ_CP017671.1 | *P. rettgeri* strain RB151 | *gne* | *gnu* | 83.23 |
| NZ_AP022375.1 | *P. rettgeri* strain BML2576 | *gne* | *gnu* | 83.44 |
| NZ_AP022372.1 | *P. rettgeri* strain BML2526 | *gne* | *gnu* | 83.44 |
| NZ_LS483422.1 | *P. heimbachae* strain NCTC12003 | *galE* | *galE_3* | 80.06 |
| NZ_LR134396.1 | *P. rustigianii* strain NCTC8113 | *galE* | *galE_3* | 83.33 |
| NZ_LR134189.1 | *P. rustigianii* strain NCTC6933 | *galE* | *galE_2* | 82.65 |
| NZ_CP077388.1 | *P. rettgeri* strain FDAARGOS 1451 | *galE* | *galE_2* | 77.94 |

**Table S6**. Summary of genes identified within the O-antigen operon homologous to those from the Ovchinnikova et al. study. Gene homolog or its product name is provided according to Prokka annotation.

| | | | | |
|---|---|---|---|---|
| NZ_CP077317.1 | *P. rettgeri* strain FDAARGOS 1450 | *galE* | *galE_1* | 71.8 |
| NZ_CP077260.1 | *P. rettgeri* strain FDAARGOS 1449 | *galE* | *galE_2* | 78.35 |
| NZ_CP076407.1 | *P. rettgeri* strain PreM15758 | *galE* | *galE_2* | 70.82 |
| NZ_CP076406.1 | *P. rettgeri* strain PreM15973 | *galE* | *galE_2* | 70.82 |
| NZ_CP076405.1 | *P. rettgeri* strain PreM15628 | *galE* | *galE_2* | 70.82 |
| NZ_CP076258.1 | *P. rettgeri* strain W986 | *galE* | *galE_3* | 78.27 |
| NZ_CP066315.1 | *P. rettgeri* strain R39 | *galE* | *galE_3* | 78.27 |
| NZ_CP066071.1 | *P. stuartii* strain FDAARGOS_1040 | *galE* | *galE_1* | 76.48 |
| NZ_CP065420.1 | *Providencia sp.* 2.29 | *galE* | *galE_2* | 76.01 |
| NZ_CP062821.1 | *P. rettgeri* strain Res13-Sevr-LER2-35 | *galE* | *galE_3* | 71.6 |
| NZ_CP060726.1 | *P. rettgeri* strain YPR25 | *galE* | *galE_2* | 78.31 |
| NZ_CP059348.1 | *P. rettgeri* strain ZG2016M39-3 | *galE* | *galE* | 75.47 |
| NZ_CP059347.1 | *P. rettgeri* strain 2353217 | *galE* | *galE_3* | 77.97 |
| NZ_CP059346.1 | *P. alcalifaciens* strain 1701003 | *galE* | *galE_2* | 97.37 |
| NZ_CP059345.1 | *P. rettgeri* strain 2055 | *galE* | *galE_3* | 73.7 |
| NZ_CP059298.1 | *P. rettgeri* strain P01 | *galE* | *galE_1* | 71.9 |
| NZ_CP058958.1 | *P. rettgeri* strain G0519 | *galE* | *galE_2* | 71.9 |
| NZ_CP053896.1 | *P. rettgeri* strain YPR31 | *galE* | *galE_2* | 78.31 |
| NZ_CP048796.1 | *P. vermicola* strain P8538 | *galE* | *galE_2* | 75.5 |
| NZ_CP048621.1 | *P. stuartii* strain MF1 | *galE* | *galE_2* | 76.48 |

**Table S6**. Summary of genes identified within the O-antigen operon homologous to those from the Ovchinnikova et al. study. Gene homolog or its product name is provided according to Prokka annotation.

| | | | | |
|---|---|---|---|---|
| NZ_CP044076.1 | *P. stuartii* strain FDAARGOS_645 | *galE* | *galE_2* | 73.78 |
| NZ_CP042861.1 | *Providencia* sp. 1709051003 | *galE* | *galE_2* | 78.07 |
| NZ_CP042860.1 | *Providencia* sp. 1701091 | *galE* | *galE* | 71.69 |
| NZ_CP042859.1 | *Providencia* sp. 1701011 | *galE* | *galE* | 71.69 |
| NZ_CP039844.1 | *P. rettgeri* strain Pr-15-2-50 | *galE* | *galE_2* | 78.07 |
| NZ_CP031123.2 | *P. huaxiensis* strain WCHPr000369 | *galE* | *galE_3* | 71.57 |
| NZ_CP029736.1 | *P. rettgeri* strain AR_0082 | *galE* | *galE* | 72.71 |
| NZ_CP028384.1 | *P. heimbachae* strain 99101 | *galE* | *galE_2* | 73.78 |
| NZ_CP027418.1 | *P. rettgeri* strain FDAARGOS_330 | *galE* | *galE_1* | 78.75 |
| NZ_CP027398.1 | *P. stuartii* strain FDAARGOS_291 | *galE* | *galE_2* | 72.56 |
| NZ_CP026704.1 | *P. stuartii* strain AR_0026 | *galE* | *galE_2* | 73.73 |
| NZ_CP023536.1 | *P. alcalifaciens* strain FDAARGOS_408 | *galE* | *galE* | 97.66 |
| NZ_CP017671.1 | *P. rettgeri* strain RB151 | *galE* | *galE_3* | 78.75 |
| NZ_CP017054.1 | *P. stuartii* strain BE2467 | *galE* | *galE_2* | 74.02 |
| NZ_CP014024.2 | *P. stuartii* strain FDAARGOS_145 | *galE* | *galE_2* | 76.48 |
| NZ_CP008920.1 | *P. stuartii* strain ATCC 33672 | *galE* | *galE_1* | 76.48 |
| NZ_AP022375.1 | *P. rettgeri* strain BML2576 | *galE* | *galE_2* | 72.86 |
| NZ_AP022374.1 | *P. stuartii* strain BML2537 | *galE* | *galE_2* | 76.41 |
| NZ_AP022373.1 | *P. rettgeri* strain BML2531 | *galE* | *galE_2* | 71.97 |

**Table S6**. Summary of genes identified within the O-antigen operon homologous to those from the Ovchinnikova et al. study. Gene homolog or its product name is provided according to Prokka annotation.

| | | | | |
|---|---|---|---|---|
| NZ_AP022372.1 | *P. rettgeri* strain BML2526 | *galE* | *galE_1* | 72.86 |
| NZ_AP022371.1 | *P. rettgeri* strain BML2496 | *galE* | *galE_2* | 70.82 |
